# Core spliceosomal Sm proteins as constituents of cytoplasmic mRNPs in plants

**DOI:** 10.1101/709550

**Authors:** Malwina Hyjek-Składanowska, Mateusz Bajczyk, Marcin Gołębiewski, Przemysław Nuc, Agnieszka Kołowerzo-Lubnau, Artur Jarmołowski, Dariusz Jan Smoliński

## Abstract

In light of recent studies, many of the cytoplasmic posttranscriptional mRNA processing steps take place in highly specialized microdomains referred to as cytoplasmic bodies. These evolutionarily conserved microdomains are sites of regulation for both mRNA translation and degradation. It has been shown that in the larch microsporocyte cytoplasm, there is a significant pool of Sm proteins not related to snRNP complexes. These Sm proteins accumulate within distinct cytoplasmic bodies (S-bodies) that also contain mRNA. Sm proteins constitute an evolutionarily ancient family of small RNA-binding proteins. In eukaryotic cells, these molecules are involved in pre-mRNA splicing. The latest research indicates that in addition to this well-known function, Sm proteins could also have an impact on mRNA at subsequent stages of its life cycle. The aim of this work was to verify the hypothesis that canonical Sm proteins are part of the cytoplasmic mRNP complex and thus function in the posttranscriptional regulation of gene expression in plants.

## INTRODUCTION

Posttranscriptional regulation of gene expression plays an essential role in all aspects of the cell life cycle, including cell development, differentiation, survival, homeostasis, adaptation to stress and response to environmental signals. Various mechanisms that enable efficient transcription, depending on actual need, have evolved in eukaryotic cells. Once mRNA is transcribed, it is bound by RNA-binding proteins (RBPs), key regulators of both mRNA stability and translation (Abdelmohsen, 2012). RBPs accompany mRNAs throughout their whole life cycle and guide the transcripts at every step of their maturation and turnover (Hieronymus and Silver, 2004; Keene, 2007). Due to the complexity and dynamic nature of mRNP complexes, determination of the full composition of a single mRNP complex remains a challenge. It has been shown that RNA binding proteins constitute 3 to 11% of bacterial and eukaryotic proteomes (Anantharaman *et al.*, 2002).

The cytoplasm plays the key role in the regulation of gene expression by controlling the level of mRNA translation and decay. The transport, localization and degradation of mRNA within this compartment are highly ordered processes. This complexity is not surprising considering the high content of mRNA in the cytoplasm (up to 150 000 molecules in mammalian cells; Halbeisen *et al*. 2008), where two major, counteracting processes of mRNA translation and decay must occur at the same time. As stated in literature, a vast majority of mRNA posttranscriptional regulation occurs within the specialized microdomains referred to as cytoplasmic bodies (Moser and Fritzler, 2010; Lavut and Raveh, 2012). These nonmembrane bound, highly dynamic structures enriched in ribonucleoproteins are evolutionarily conserved, as their occurrence has been confirmed in yeasts (Sheth and Parker, 2003; Lavut and Raveh, 2012), protozoans (López-Rosas *et al.*, 2012), nematodes (Barbee *et al.*, 2002; Gallo *et al.*, 2008), insects (Liu and Gall, 2007; Lee *et al.*, 2009), amphibians (Bilinski *et al.*, 2004), mammals (Chuma *et al.*, 2003; Sen and Blau, 2005), and plants Xu and Chua, 2009; Smoliński *et al.*, 2011; Perea-Resa *i in.*, 2012). Several cytoplasmic bodies have been described to date, including P-bodies, stress granules, neuronal granules, germinal granules and nuage, among others.

The core cytoplasmic bodies in eukaryotic cells are P-bodies, which are sites of mRNA regulation, including 5’–3’ deadenylation-dependent degradation (Sheth and Parker, 2003; Beggs, 2005), miRNA-mediated decay (Sen and Blau, 2005) and mRNA stabilization and sequestration (Xu and Chua, 2009), as well as nonsense-mediated decay (NMD; Unterholzner and Izaurralde, 2004) and translational regulation (Andrei *et al.*, 2005; Kedersha *et al.*, 2005; Wilczynska *et al.*, 2005; Chu and Rana, 2006). Due to their composition, including proteins involved in both mRNA stabilization and turnover, P-bodies constitute centers of mRNA sorting where transcripts are directed to either translation or degradation (Lavut and Raveh, 2012). It has been revealed that the components of P-bodies accumulate transiently and can shuttle between the cytoplasm and other cytoplasmic granules (Kedersha *et al.*, 2005; Wilczynska *et al.*, 2005; Moser and Fritzler, 2010). Moreover, there is evidence that molecules such as GW182 protein, AGO2-miRNP and mRNA itself can be loaded into specific microvesicles, referred to as extracellular exosomes, and transferred between cells in fully functional form (Gibbings *et al.*, 2009). Thus, the mRNA life cycle involves constant dynamic changes in mRNP composition and subcellular localization, where mRNPs can shuttle between polysomes and cytoplasmic bodies, depending on external stimuli and the current metabolic needs of the cell. Despite growing research on the posttranscriptional regulation of mRNA utilization and its spatial organization within the cytoplasm, the full mechanisms and functional roles of mRNP assembly into higher order microdomains need to be elucidated.

Here, we show that canonical core spliceosomal proteins (Sm proteins) constitute a part of the cytoplasmic mRNP complex in European larch (*Larix decidua* L.) microsporocytes. These cytoplasmic bodies were localized after transcriptional bursts, indicating that these microdomains are formed as a result of pulsed, extensive mRNA synthesis. The European larch microsporocytes are male germline precursors that have been shown to be highly active periodically during diplotene (Kołowerzo-Lubnau *et al*., 2015). The diplotene stage lasts approximately 5 months and it is possible to divide it into several substages and to observe each of them in details. It has been demonstrated that in *Larix* species, the chromatin is subject to substantial rearrangements, and the microsporocyte is highly metabolically active, increasing its volume 6-fold during this stage (Zhang *et al*., 2008; Kołowerzo-Lubnau *et al*., 2015). In *Larix decidua*, metabolic activity is strictly driven by cyclic chromatin contraction and diffusion. Five periods of chromatin relaxation (diffuse stages) separated by four periods of genome contraction are distinguished during diplotene. In many animal and plant species, the diplotene stage is characterized by significant chromatin relaxation, referred to as the diffuse stage of meiosis (Klasterska, 1976; Cenci *et al*., 1994; Sheehan and Pawlowski, 2009; Comizzoli *et al*. 2011, She *et al*., 2013; Colas *et al*., 2017;). As a result, RNA synthesis is not a continuous process but occurs in transcriptional bursts — the diffuse stages are accompanied by large increases of *de novo* transcription of both mRNAs and rRNAs, as well as the expression and assembly of splicing subunits (Smoliński *et al*., 2007, 2011; Hyjek *et al*., 2015). Here, we show that in the larch microsporocyte during diplotene after extensive mRNA synthesis in the diffuse stage Sm proteins accumulate in the cytoplasm within distinct cytoplasmic bodies that also contain mRNA.

Sm proteins constitute an evolutionarily ancient family of small RNA-binding proteins. In eukaryotic cells, these molecules are primarily involved in pre-mRNA splicing. Only a few studies have indicated that, in addition to this well-known function, Sm proteins could also impact mRNA at subsequent stages of its life cycle. In our research, using larch microsporocytes as a model system, we show that there is a significant pool of Sm proteins accumulated within distinct cytoplasmic bodies that also contain polyadenylated RNA (poly(A) RNA). A correlation between the cyclic occurrence of the cytoplasmic bodies and the metabolic activity of the cell was shown. A total of 222 mRNAs were identified as cytoplasmic partners for Sm proteins, and they comprise three major categories of Sm protein-associated mRNAs that code for proteins related to (i) ribosomes/translation, (ii) mitochondria/energy metabolism and (iii) chloroplasts/photosynthesis. Moreover, the distribution analysis of P-body or stress granules markers (i.e., LSm4, DCP2, AGO1 and RS28) showed that Sm protein-mRNA cytoplasmic bodies are not functionally related to these bodies.

Based on the results obtained in this work and studies on other model eukaryotic cells, we propose that Sm protein-mRNA bodies constitute newly described cytoplasmic domains, as implicated in the posttranscriptional regulation of highly expressed transcripts, particularly in cells in which mRNA synthesis occurs in transcriptional bursts.

The aim of this work was to verify the hypothesis that canonical Sm proteins are part of the cytoplasmic mRNP complex and thus function in the posttranscriptional regulation of gene expression in plants.

## RESULTS

### S-bodies: cytoplasmic bodies rich in the Sm proteins and polyadenylated mRNA

A double labeling assay of the core spliceosomal Sm proteins and poly(A) RNA revealed a remarkable distribution of these molecules within the cytoplasm (Figure 1, Figure S1).

**Figure 1.**
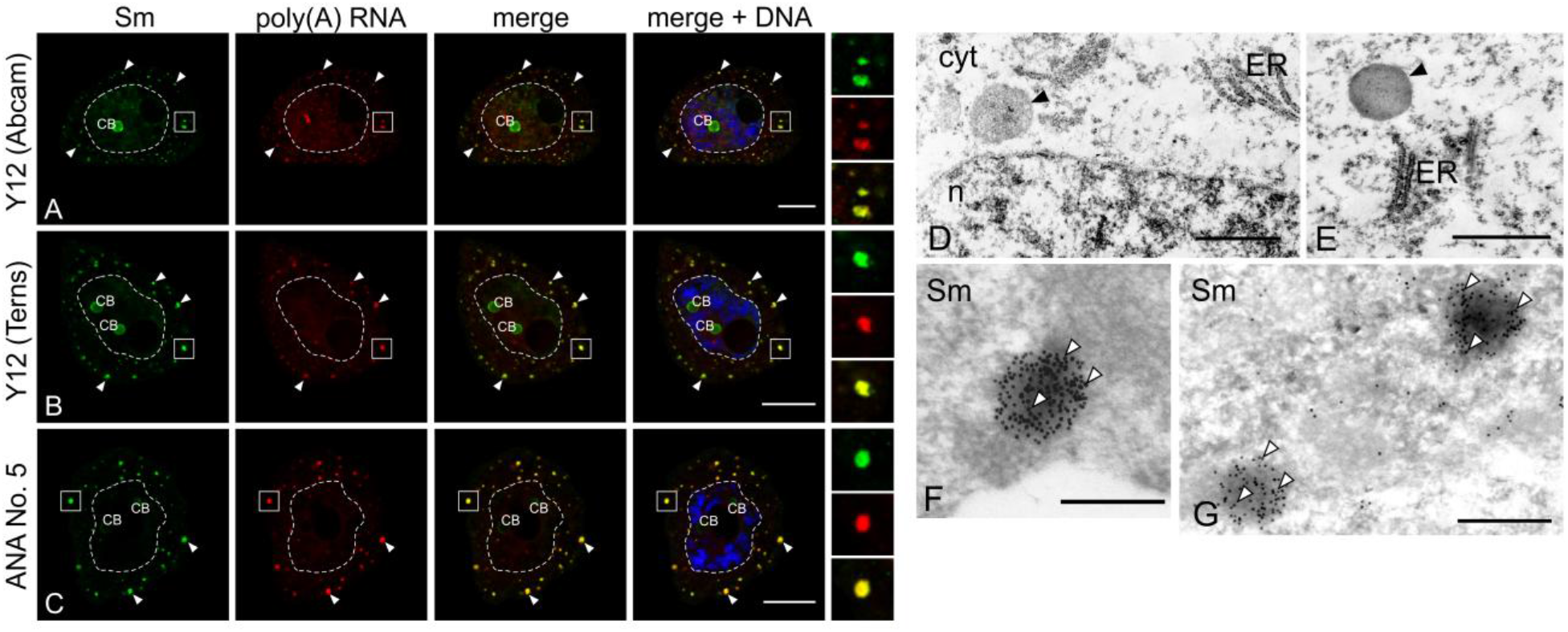
S-bodies: cytoplasmic bodies rich in the Sm proteins and poly(A) RNA. **A-C**, Localization of Sm proteins and poly(A) RNA in microsporocytes during diplotene. Visible numerous accumulation of Sm proteins in the colocalization of poly(A) RNA (arrowheads). The right panel represents the magnification of the cytoplasm, which is marked with a square. CB – Cajal body. Bar −15 μm. **D-E**, Ultrastructural analysis of microsporocytes. Visible nonmembrane bound cytoplasmic structures (arrowheads).n - nucleus, cyt - cytoplasm, ER - endoplasmic reticulum. Bar - 1 μm. **F-G**, Location of the Sm proteins in the cytoplasm of the microsporocytes as determined by the immunogold method. Visible accumulations of gold grains (20 nm) in isolated cytoplasmic clusters (arrowheads). Bar - 0.5 μm.

Distinct 0.5 - 1 µm cytoplasmic foci were observed, in which a significant amount of polyadenylated transcripts had accumulated, in colocalization with Sm proteins. To verify the specificity of the observed labeling, three different antibodies were used for Sm protein localization, namely, commercially available Y12 MAbs (Figure 1a), Y12 MAbs from Matera’s laboratory (Figure 1b) and the ANA No. 5 human reference anti-serum (Figure 1c). All the antibodies used showed a similar pattern of labeling with a well-established, characteristic nuclear pool of Sm proteins present in the nucleoplasm and Cajal bodies (CBs). Moreover, numerous discrete foci of Sm protein accumulation were observed in the cytoplasm and colocalized with poly(A) RNA (Figure S1). The ultrastructural analysis revealed spherical, nonmembrane bound microdomains with a diameter of 0.4–1 µm (Figure 1d,e). The immunogold assay confirmed that a noticeable portion of the Sm proteins accumulated within distinct cytoplasmic structures (Figure 1f,g). Based on these observations and taking into consideration the general nomenclature used for this type of microdomains, we propose a name of S-bodies for these structures.

### S-bodies are formed during the cytoplasmic export of poly(A) RNA

Larch microsporocytes exhibit a rigorously regulated pattern of metabolic activity. During the diplotene stage of meiosis, five bursts of *de novo* synthesis of poly(A) RNA occur, separated by periods of transcriptional silencing (Kołowerzo-Lubnau *et al*., 2015). Each of these cellular poly(A) RNA turnover cycles comprises subsequent stages of nuclear synthesis, cytoplasmic export and degradation of mRNA (Figure 2). A detailed microscopic analysis showed that the S-bodies emerge periodically in each of the five cycles of poly(A) RNA turnover. Based on the transcription rate and localization of the poly(A) transcripts, the longest and most intensive fourth cycle could be divided into seven stages. Shortly after intensive synthesis in the nucleus (stages I and II, Figure 2a), poly(A) RNAs are transported to the cytoplasm, as indicated by an increasing level of labeling in this compartment (stages III through VII; Figure 2a). A remarkable ring-like accumulation of poly(A) RNA was observed around the nucleus, reflecting the pool of newly exported transcripts. Additionally, starting at stage III of the cycle, distinct foci of the poly(A) RNA and Sm proteins were observed and represented the S-bodies. At stage IV, the number of bodies was the highest in the whole cycle (380 ± 35 bodies per cell, Figure 2b). During the subsequent stages of cytoplasmic RNA localization, this number was consequently decreased (stages V through VII, Figure 2b). It should be emphasized, however, that throughout the whole period when S-bodies were observed, there was a significant colocalization of both poly(A) RNA and Sm proteins (PCC = 0.69 ± 0.02, Figure 2d) in the cytoplasmic pool. Furthermore, the Manders’ overlap calculation demonstrated that a substantial fraction of Sm proteins in the cytoplasmic pool had colocalized with poly(A) RNA (67,43 ± 2,90%, Figure 2e).

**Figure 2:**
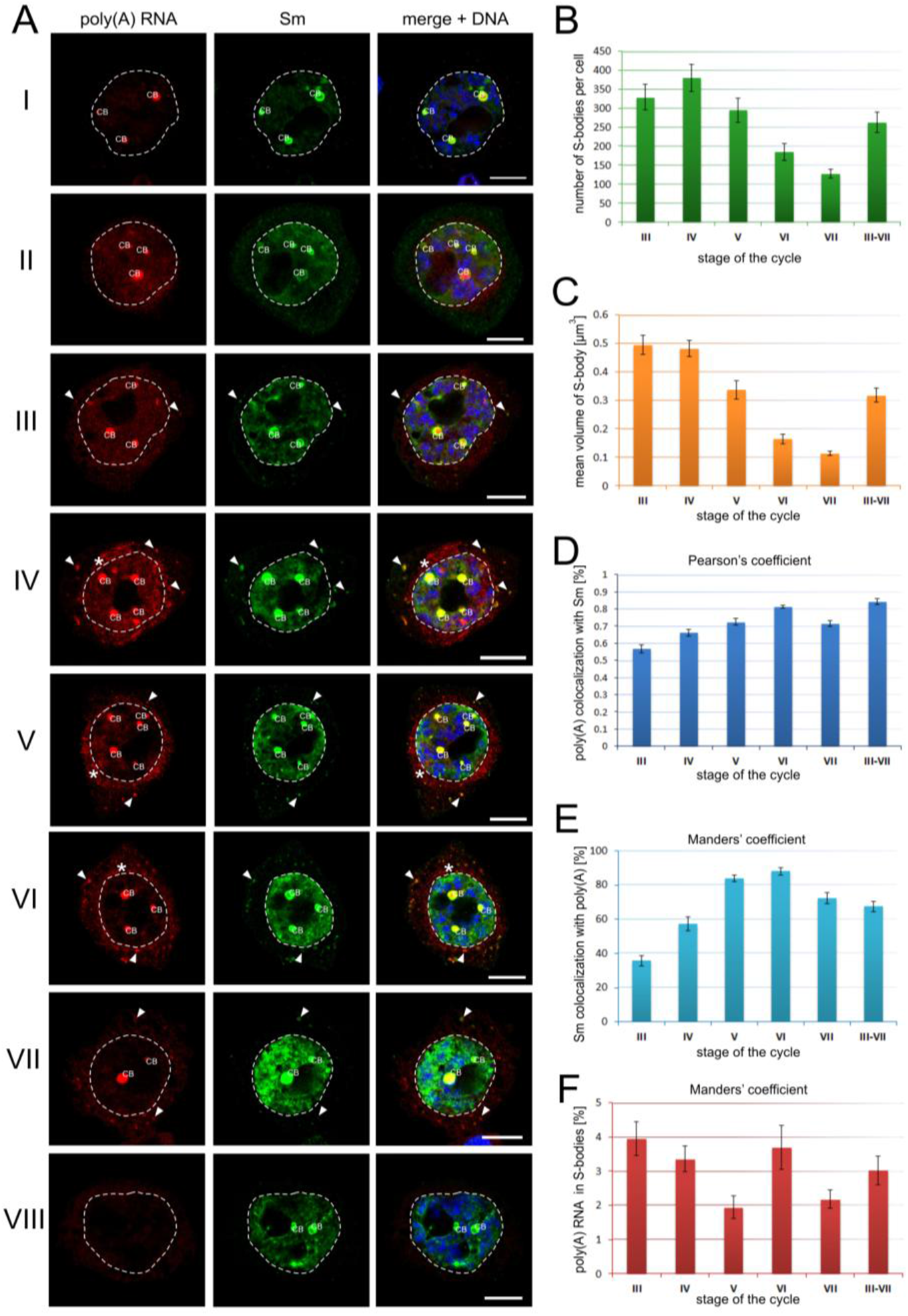
The longest and most intensive fourth cycle of poly(A) RNA turnover during diplotene stage of the first meiotic prophase. **A**, The spatial and temporal distribution of the Sm proteins in the fourth cycle of poly(A) RNA turnover. I - VIII - successive stages of the cycles of poly(A) RNA turnover. A detailed description is in the text. CB - Cajal body. Bar −15 μm. **B**, The mean number of S-bodies per cell in subsequent stages of the cycles of poly(A) RNA turnover. Error bars indicate the standard error of the mean (SEM). p <0.05. **C**, The mean volume of S-bodies in subsequent stages of the cycles of poly(A) RNA turnover. **D-E**,Analysis of the colocalization of poly(A) RNA and Sm proteins in the cytoplasm at subsequent stages of the poly(A) RNA cycle in the cell. The values are shown as the mean ± standard error of the mean (SEM). PC - Pearson colocalization coefficient. **F**, The percentages were calculated on the basis of the Manders’ colocation coefficient, in which the value of 1 was converted to 100% colocalization. Error bars indicate the standard error of the mean (SEM). a-b p <0.01, a-c and b-c p <0.001.

The overlap coefficient gradually increased as the cytoplasmic export of mRNAs proceeded: from 35.76 ± 3.09% at stage III to 88.03 ± 2.13% of the total cytoplasmic Sm protein pool at stage VI (Figure 2e). We also estimated the percentage of the cytoplasmic poly(A) RNA pool that was in S-bodies during the fourth cycle of transcriptional activity at the diplotene stage. The analysis revealed that 3.08 ± 0.42% of the total cytoplasmic pool of poly(A) RNA had accumulated in cytoplasmic bodies, and this accumulation was consistent throughout the cytoplasmic phase of the RNA cell cycle (differences between stages were not statistically significant, ANOVA, p>0.05, Figure 2f).

To test the relationship between the occurrence of cytoplasmic bodies and the metabolic state of the cell, a quantitative analysis of the marker for transcriptional activity (active RNA pol II) was performed in the 3 cycles of the most intensive poly(A) RNA turnover (Figure 3). During the stages of RNA cytoplasmic export (stages III-VII, see Figure 2a), i.e., when the Sm proteins and polyadenylated transcripts accumulate within cytoplasmic bodies, the transcriptional activity was considerably decreased compared to the transcriptional activity in the nuclear stages of mRNA synthesis (stages I-II, see Figure 2a; Figure 3a). In contrast, the level of the marker for potential translational activity (5S rRNA) showed no correlation (Figure 3b). These results indicate that the formation of S-bodies in the cytoplasm is slightly postponed with respect to bursts of nuclear mRNA synthesis and that these microdomains appear in periods of decreased transcriptional activity.

**Figure 3:**
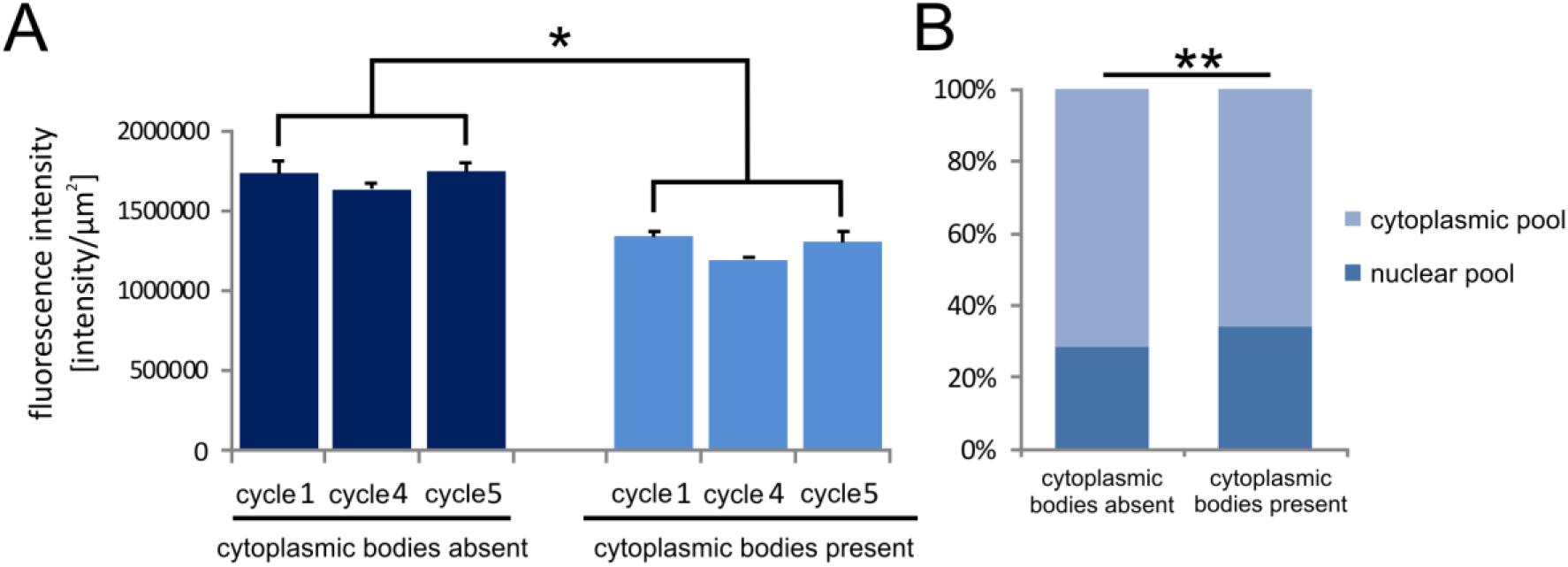
Quantitative analysis of the markers for transcriptional activity (RNA pol II and 5S rRNA). **A**, Quantitative analysis of the active level of RNA polymerase II in the different cycles of poly(A) RNA synthesis in the presence and absence of Sm protein/poly(A) RNA bodies in the cytoplasm of the cell. The error bars indicate the standard error of the mean (SEM). * p <0.05. **B**, analysis of the nucleus:cytoplasm ratio of 5S rRNA in cells containing and not containing Sm protein/poly (A) RNA bodies. ** p <0.01.

### S-bodies are not sites of spliceosomal particle accumulation

The well-known and broadly documented role of Sm proteins in the nucleus is their participation in pre-mRNA splicing. Sm proteins bind uridine-rich small nuclear RNAs (U snRNAs) in the form of a heptameric ring around the transcript, creating the core spliceosomal ribonucleoparticles U1, U2, U4 and U5 snRNPs (Zieve *et al*., 1988; Baserga and Steitz, 1993; Raker *et al*., 1999, Shaw *et al*., 2008). The assembly of U snRNP involves a highly dynamic cytoplasmic stage at which the Sm proteins are loaded onto the U snRNA transcripts and the 5’ trimethyl-guanosine (m3G) cap is formed (Mattaj, 1986; Fischer *et al*., 1997; Urlaub *et al*., 2001; Yong *et al*., 2010). Moreover, our previous research (Hyjek *et al*., 2015) revealed that, during larch meiosis, the cytoplasmic stage of U snRNP assembly can occur in two spatially distinct manners: it may be diffused throughout the cytoplasm or localized within snRNP-rich cytoplasmic bodies (CsBs). The assembly mode depends on the rate of U snRNP *de novo* formation and the amount of U snRNP in the nucleus. Thus, it could not be precluded that the S-bodies are related to the assembly and/or sequestration of spliceosomal elements.

To test this possibility, the cellular distribution of both the nucleotide (U1, U2, U4, U5 snRNA, and m3G cap) and protein (U2B’’) components of the U snRNPs were analyzed in relation to the S-bodies (Figure 4, Figure S2). All the spliceosomal components were specifically localized within the nucleus, with significant accumulation in Cajal bodies (CBs). In contrast, no signal in the form of distinct clusters of U snRNA, m3G cap or U2B’’ protein was observed in the cytoplasm, indicating that the formation of S-bodies does not coincide with the stage of U snRNP assembly in the cytoplasm. The lack of any U snRNP components and the presence Sm proteins in the S-bodies demonstrates that the bodies represent the cytoplasmic pool of canonical Sm proteins acting outside of the spliceosome, and that this localization is presumably related to mRNA.

**Figure 4:**
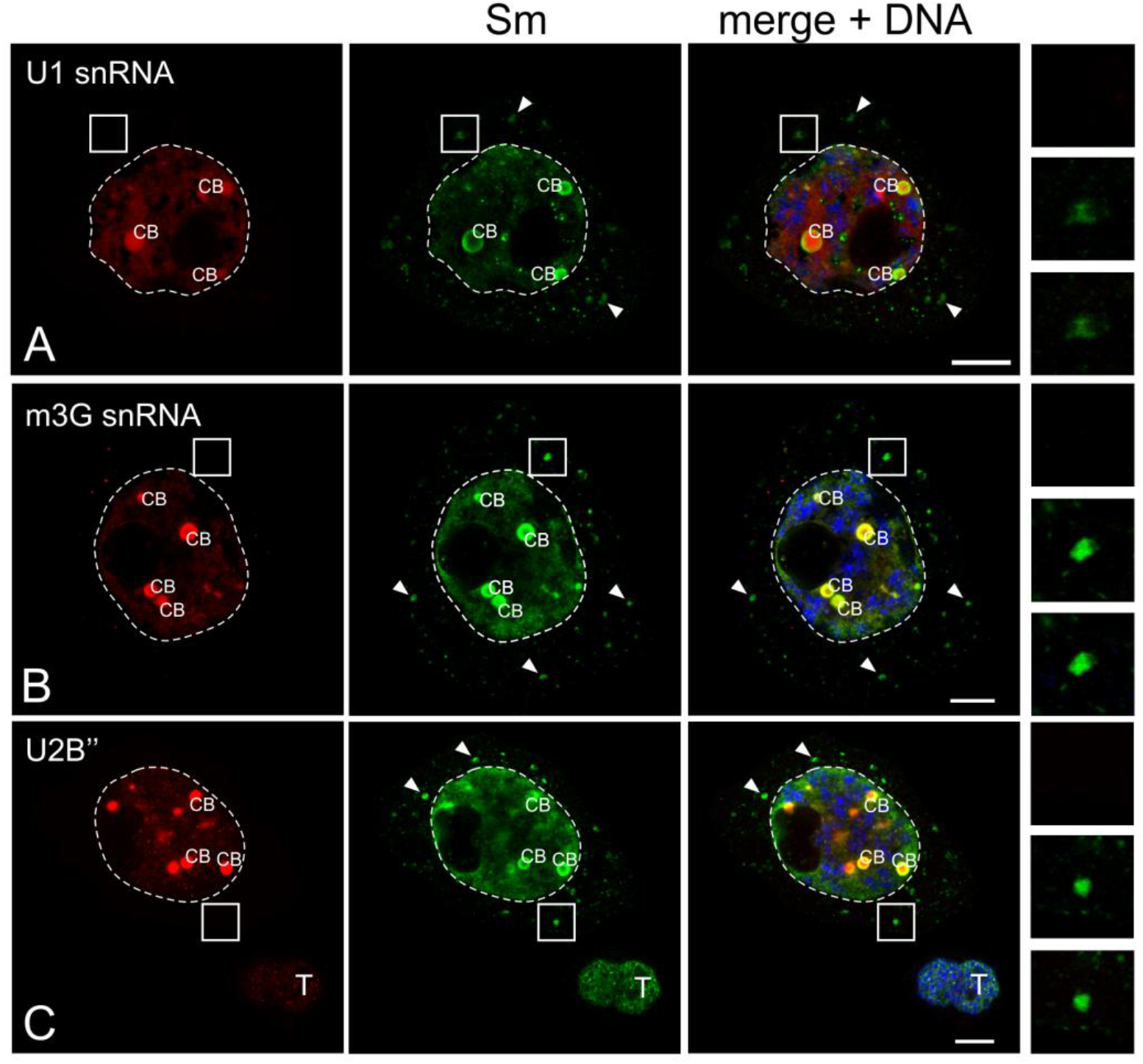
S-bodies are not sites of the U snRNPs assembly. Analysis of the Sm protein distribution in the colocalization with selected splicing elements. There was no accumulation of U1 snRNA (**A**), m3G cap (**B**) or U2B” proteins (**C**) in the S-bodies (arrowheads). The right panel represents the magnification of the cytoplasm, which is marked with a square. CB - Cajal body. T - anther tapetum cell. Scale of 10 μm.

### Identification of Sm protein-associated cytoplasmic mRNA transcripts

To identify the mRNA transcripts that accumulate within S-bodies, we employed an RNA-immunoprecipitation (RIP) approach against the Sm proteins from the larch microsporocyte cytoplasm, followed by high-throughput sequencing of the immunopurified RNAs (RIP-seq).

Since all three anti-Sm antibodies used for the assays *in situ* could label the cytoplasmic bodies with comparable specificity and strength (see Figure 1), an additional validation of the results was performed. All tested immunoglobulins specifically precipitated the SmB protein, however only Y12 antibody (not commercial), also effectively precipitated the low-molecular SmD1-SmD3 proteins (Figure S3). In order to confirm the specificity of the antibodies used for the isolation and detection of Sm proteins in Larix decidua, identification of proteins after immunoprecipitation (IP) was performed using these antibodies by mass spectrometry (IP-SDS-PAGE-MS method). On the basis of the results obtained, in conjunction with the predicted mass of the studied polypeptides, seven of the eight core Sm proteins were identified: SmF (11 kDa), SmE (12 kDa), SmD1 (13 kDa), SmD2 (14 kDa), SmD3 (16 kDa), SmB (25 kDa) and SmB’ (26 kDa) (Figure S4).

Finally, an additional validation of the results from the RNA immunoprecipitation analyses was performed to select the most efficient antibody to use for further analysis (Figure 5a). The enrichment of U2 snRNA — a canonical cellular partner of the Sm proteins was estimated, and the results were normalized to 5S rRNA (which does not interact with Sm proteins). The experiment revealed that the Y12 MAbs obtained from Matera’s laboratory was the most effective for the RIP assay, precipitating over 1130 times more U2 snRNA than the control (mock IP; no antibody added). This was significantly different than the 84-fold and 12-fold enrichment obtained from using ANA No. 5 and the commercially available Y12, respectively.

**Figure 5:**
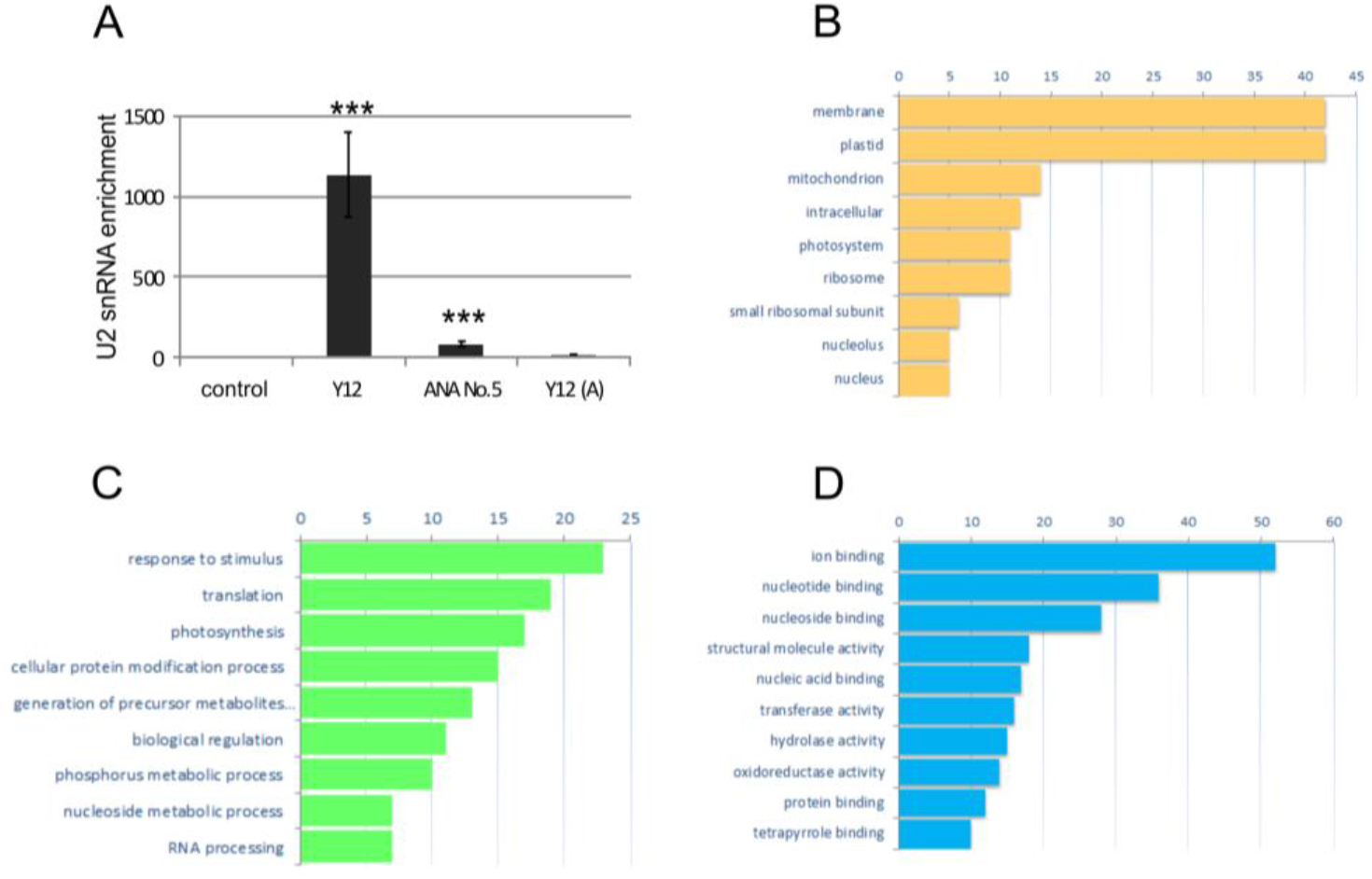
Identification of mRNA transcripts. **A**, Determination of the efficiency of U2 snRNA immunoprecipitation using different anti-Sm protein antibodies (Y12, Ana No. 5, and Y12 (A)) by real-time (RT)-PCR. The graph shows the enrichment values relative to the control (for the control, the value was 1), normalized to the level of 5S rRNA. The error bars indicate the standard error of the mean (SEM). *** p <0.001. D-F The functional annotation of cytoplasmic mRNA precipitating with Sm proteins: biological process (**B**), molecular function (**C**), and cell compartment (**D**). Numbers in the horizontal line indicate number of genes, enrichment ratios to control were significant for all RIP transcripts.

The Sm protein-associated transcripts are listed in Supplementary Data 1. A total of 245 transcripts were identified as cytoplasmic partners of Sm proteins, of which 222 were classified as mRNAs. Most of the transcripts were homologous to *Pinaceae*. The Gene Ontology (GO) functional annotation revealed that the cytoplasmic Sm interactome consisted of mRNAs encoding a vast range of proteins, localizing to many cell compartments and linking a number of metabolic processes. A total of 141 sequences (58%) were annotated with 728 GO terms. This functional analysis revealed three major categories of Sm protein-associated mRNAs, and they coded for proteins related to (i) ribosomes/translation, (ii) mitochondria/energy metabolism and (iii) chloroplasts/photosynthesis (Figure 5b-d).

To test whether the immunoprecipitated mRNAs are indeed accumulated within S-bodies, the spatial distribution of the transcripts was analyzed *in situ*. From the 222 Sm protein-associated mRNAs 16 were selected, representing different cellular functions and localization. A double labeling assay of the larch microsporocytes was performed for the Sm proteins and each of the selected mRNAs (Figure 6, Figure S5).

**Figure 6:**
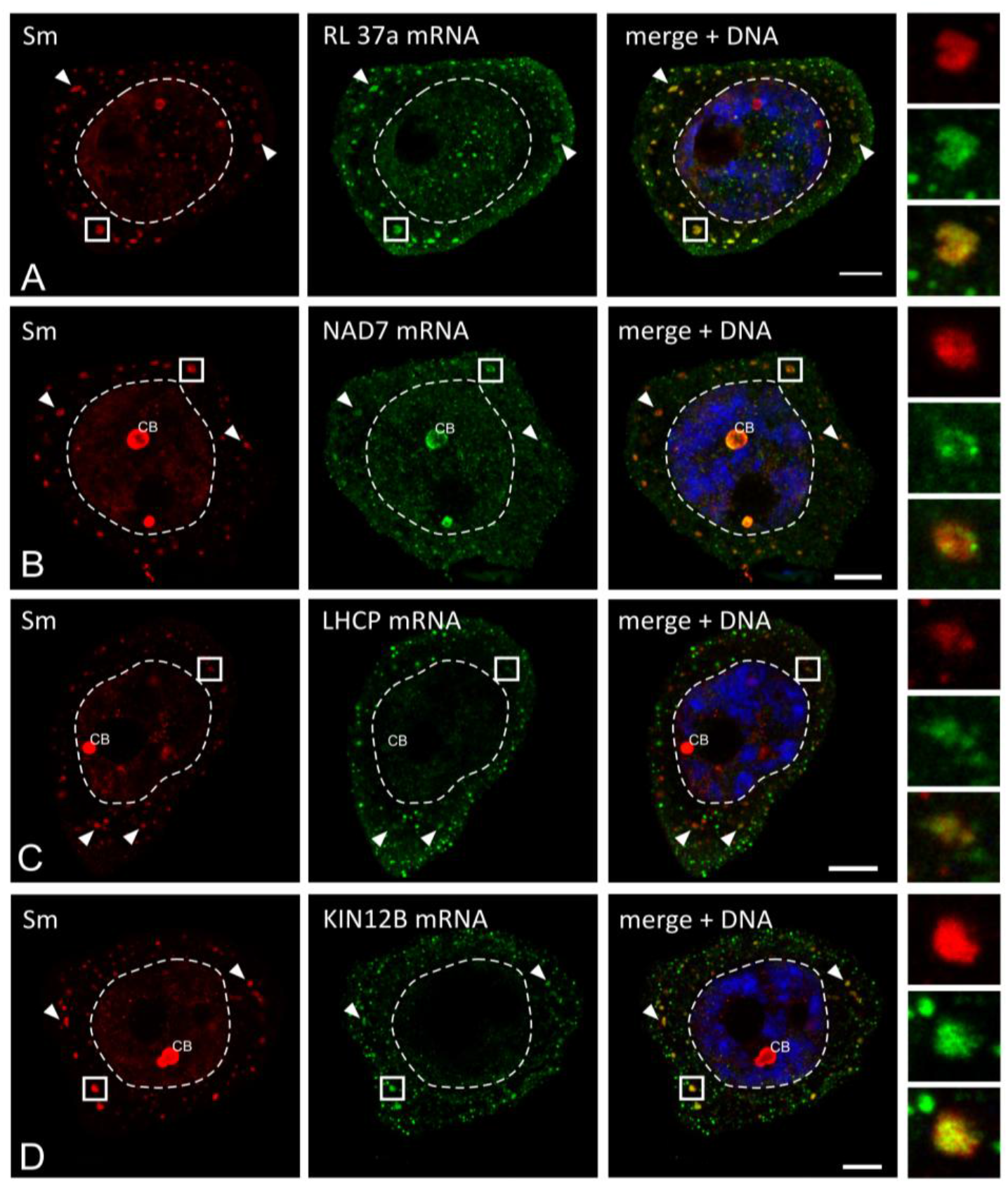
*In situ* localization of mRNA precipitating with Sm proteins. The results represent the stages of the IV - VI poly(A) RNA turnover. Numerous clusters of mRNA that colocalized with the Sm proteins (arrowheads) are visible in the cytoplasm. The right panel represents the magnification of the cytoplasm, which is marked with a square. **A**, RL 37a - large ribosome subunit protein 37a; **B**, NAD7 - NADH 7 dehydrogenase subunit; **C** - LHCP - protein binding chlorophyll a-b; **D**, KIN12b - kinesin 12b-like protein, CB - Cajal body. Bar −10 μm.

The microscopic analysis confirmed the specific locations of the 13 mRNAs within the cytoplasmic bodies. This specific distribution pattern was observed for mRNAs that encoded diverse proteins, including those involved in translation (Figure 6a, Figure S5a,c,d), energy metabolism (Figure 6b, Figure S5f) and photosynthesis (Figure 6c, Figure S5i), as well as structural proteins of the cytoskeleton (Figure 6d, Figure 5Sb). Additionally, S-bodies were enriched in several mRNAs related to protein folding (Figure 5Sg,h), proteasomal degradation (Figure 5Sj) and miRNA biogenesis (Figure 5Se). A different pattern of localization was observed for only 3 mRNAs (Figure S5k-m). Transcripts encoding cellulase (Figure S5k) and tubulin (Figure 5Sl) showed a diffuse pattern of cytoplasmic distribution, with no distinct concentration in any particular area.

Double labeling of these mRNAs with Sm proteins revealed that both of them were found in the cytoplasmic bodies, although without noticeable accumulation (i.e., there were no differences between the level of staining within the cytoplasmic bodies and in the surrounding cytoplasm). For the mRNA that encodes a light-independent protochlorophyllide oxidoreductase (LI-POR), the level of staining was below the limit of detection (Figure 5Sm).

In the next step, we performed an additional analysis of the SmD mRNAs that were identified in the cytoplasmic transcriptome of the larch microsporocytes, but not enriched in the Sm protein-immunoprecipitated pool of transcripts obtained via RIP-seq assay. The lack of cytoplasmic accumulation within these microdomains was observed in the case of mRNAs encoding SmD1, SmD2 and SmE proteins (Figure S6). This finding demonstrates that the S-bodies are not sites of *de novo* Sm protein synthesis.

### The cytoplasmic Sm proteins interactome

The next aim was to examine whether the S-bodies are functionally related to other microdomains, e.g., P-bodies and/or stress granules. To select potential markers for these structures, the cytoplasmic Sm protein interactome was identified. The cytoplasmic fraction of the larch anther cells was subjected to anti-Sm immunoprecipitation, which was subsequently assessed by mass spectrometry. A total of 118 proteins were detected (Table S1, S2). Among the potential markers for the cytoplasmic bodies, LSm2, LSm3 and LSm4 were identified for P-bodies, and the ribosomal proteins eIF2/IF5, RL10e, RS1, RS3 and RS28 were characteristic for the stress granules (Table S2).

Similar to the Sm protein RNA interactome, a significant fraction of the proteic interactome included proteins related to (i) ribosomes/translation (eIF2/IF5, RL10e, RS1, RS3, RS28, and OVA2), (ii) mitochondria/energy metabolism (SHM1 and SHM2 methyltransferases, IDH dehydrogenases, and ATM1 ABC transporters) and (iii) plastids/photosynthesis (RCA RuBisCO activase and TKL1 transketolase) (Table S1, S2). Furthermore, two RNA-binding protein families were identified as coprecipitating with Sm proteins: Alba (A*cetylation Lowers Binding Affinity*) and ECT (*evolutionarily conserved C-terminal region*). Alba refers to a broad group of RNA- and DNA-binding proteins implicated in chromatin organization, regulation of transcription and translation, and RNA metabolism (Goyal *et al.*, 2016). It has been shown that these proteins are involved in the regulation of plant cell development where they perform essential roles in the sperm cell specification in *A. thaliana* (Borg *et al.*, 2011). Fewer ECT proteins have been investigated compared to mRNA-binding proteins. The scarce ECT research indicates the role of ECT1 and ECT2 in calcium signal transduction from the cytoplasm to the nucleus that affects gene expression (Ok *et al.*, 2005). Based on the IP-MS analysis and taking into consideration the availability of specific antibodies, three markers of protein degradation (LSm4, DCP2, and AGO1) and one translation regulator (RS28) were selected for the determination of *in situ* localization. All the proteins were distributed throughout the microsporocyte cytoplasm, with RS28 accumulated additionally in the nucleolus (Figure 7). None of the investigated proteins colocalized with the S-bodies, indicating that these cytoplasmic bodies are not sites of mRNA degradation and are not involved in translational repression. Furthermore, no distinct foci for LSm4 (Figure 7a-d), DCP2 (Figure 7e-h), AGO1 (Figure 7i-n) and RS28 (Figure 7o-s) were detected in the cytoplasm, suggesting the lack of P-bodies or stress granules at this stage of meiosis in larch microsporocytes.

**Figure 7:**
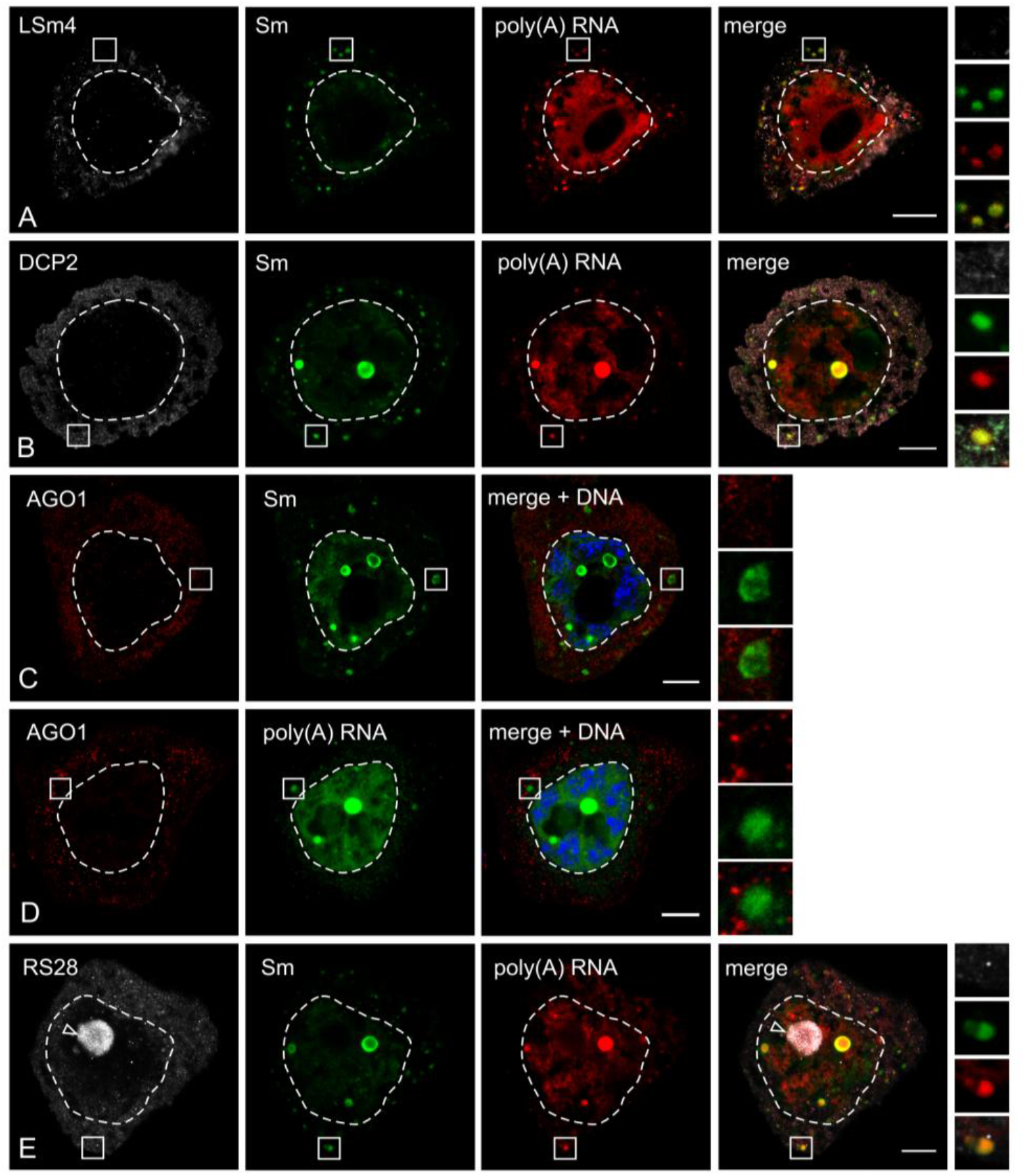
S-bodies are not functionally related to other cytoplasmic microdomains, e.g., P-bodies or stress granules. Localization of proteins related to degradation (**A**- **D**) and translation regulation (**E**) in *Larix decidua* microsporocytes. The results represent the stages of the IV - VI poly(A) RNA turnover. The right panel represents the magnification of the cytoplasm, which is marked with a square. Visible diffused fluorescence in the cytoplasm, without colocalization of the tested antigens for S-bodies and poly(A) RNA. Bar - 10 μm.

## DISCUSSION

### S-bodies — novel cytoplasmic mRNA domains in plant cells

Thus far, no distinct Sm proteins and mRNA accumulation has been observed in the cytoplasm of plant cells. However, a growing body of evidence shows a specific localization of Sm proteins and particular mRNAs in animal germ granules, which has implications for transcript localization, translational control and mRNA turnover. It was demonstrated that SmE and SmG, localizing to P-granules in *Caenorhabditis. elegans*, are involved in transcriptional quiescence in germline precursors, and that this coordination of germline differentiation is splicing-independent (Barbee *et al*.; 2002; Barbee and Evans, 2006). P-granules are structurally and functionally related to nuages in *Xenopus* oocytes, where the Sm proteins, but not other splicing elements, are localized. It was proposed that Sm proteins facilitate the transport of the mRNAs specific to germ granules (e.g. Xcat2) from the nucleus to the cytoplasm (Bilinski *et al*., 2004). In mouse spermatocytes, Sm proteins accumulate in the RNP-rich chromatoid body (Biggiogera *et al*., 1990; Moussa *et al*., 1994) and forms a complex with a marker protein of these domains - MTR1 (Chuma *et al*., 2003). In *Drosophila melanogaster*, the accumulation of SmB and SmD3 at the posterior of the developing oocyte is directly linked to the unique localization and polarization of *oskar* mRNA. The proper localization of this transcript is crucial for germ cell specification during early embryogenesis (Gonsalvez *et al*., 2010; Jaglarz *et al*. 2011). Germline and early embryonic cells are highly metabolically active, with a spatiotemporally ordered pattern of expression for distinct groups of developmental genes. The European larch microsporocytes are male germline precursors that have been shown to be highly active periodically during diplotene (Kołowerzo-Lubnau *et al*., 2015). In this context, the S-bodies are similar to the germ granules observed in animals and might be considered, to some extent, as their plant counterparts.

### Sm proteins as a part of the cytoplasmic mRNP complex

A growing body of evidence suggests additional roles for the canonical Sm proteins outside of the spliceosome in the processing, localization and translational control of mRNPs. However, little is known about which mRNPs in the pool of all transcripts are regulated by Sm proteins and what is the mechanism of their interaction. To determine which mRNAs are accumulated within the S-bodies, we performed a transcriptomic analysis of cytoplasmic Sm protein-mRNP complexes, followed by *in situ* detection analysis. The functional annotation showed that three major classes of Sm protein-associated mRNAs could be distinguished, namely, those encoding proteins related to (i) ribosomes/translation, (ii) mitochondria/energy metabolism and (iii) chloroplasts/photosynthesis. This finding is in agreement with the results obtained for *Drosophila* ovaries and HeLa cells. Lu *et al*. (2014) monstrated that among 72 Sm protein-associated, fully spliced, polyadenylated mRNAs for *Drosophila* and 30 for HeLa cells, a significant portion encoded ribosomal and mitochondrial proteins. Some of the mRNAs show an overlap in all the Sm protein interactomes studied to date — plant, insect and mammalian, including those that encode small and large ribosomal subunit proteins, translation initiation factor eIF2Bα, NADH dehydrogenase and cytochrome oxidases. Furthermore, Sm proteins were also associated with the mRNA products of intronless genes (e.g., histone H2A mRNAs in human cells, 60S ribosomal L37 mRNA in larch cells), proving that this interaction is completely independent of pre-mRNA splicing. However, the results from the animal cells do not clarify whether the Sm protein-associated mRNAs are localized to the nucleus or the cytoplasm, as the RIP-seq was performed on whole cell extracts. Nevertheless, together with the above-mentioned reports, our results indicate that the splicing-independent Sm proteins association with mRNA is conserved in invertebrates, vertebrates and plants.

Mutual posttranscriptional coordination of mRNA subpopulations has been previously described by Keene and Tenenbaum (2002). The following hypothesis was proposed based on eukaryotic posttranscriptional regulons (or operons): structurally and functionally related genes are regulated in groups by specific RNA-binding proteins to facilitate the organization of gene expression during cell growth and development (Keene and Tenenbaum, 2002; Keene, 2007). The most acknowledged RBPs involved in this type of regulation are Pumilio proteins (PUF, Puf) that participate in the posttranscriptional silencing of particular groups of genes in yeasts (Gerber *et al*., 2004), *Drosophila* (Gerber *et al*., 2006), mammals (Bohn *et al*. 2018; Goldstrohm *et al*. 2018) and plants (Tam *et al*., 2010). The posttranscriptional regulons also operate in a number of developmental processes and stress responses (Gorospe, 2003; Mazan-Mamczarz et al., 2003; Lü et al., 2006), including the immunological response in mammals (Lai *et al*., 1999; Ule *et al*., 2003; Cheadle *et al*., 2005; Lykke-Andersen and Wagner, 2005). In HeLa cells, the spliceosomal proteins U2AF (U2 auxiliary factor) and PTB (polypyrimidine tract binding protein) bind distinct groups of mature mRNAs in both the nucleus and the cytoplasm (Gama-Carvalho *et al*., 2006). While U2AF interacts predominantly with transcriptional factors and cell cycle regulator mRNAs, PTB bound transcripts encode proteins related to intracellular and extracellular trafficking and apoptosis. Moreover, SR (*Ser–Arg-*rich) spliceosomal proteins are linked to cytoplasmic export of intronless histone mRNAs (Huang and Steitz, 2001; Änkö *et al.*, 2012), stimulation of translation (Sanford *et al.*, 2004) and cytoplasmic degradation of particular transcripts (Lemaire *et al.*, 2002). Recently, the involvement of SmD1 in posttranscriptional gene silencing (PTGS) was revealed in *A. thaliana* (Elvira-Matelot *et al*. 2016). Based on the results from a genetic screening of PTGS-defective mutants, it was proposed that, in addition to its role in pre-mRNA splicing, SmD1 facilities cytoplasmic PTGS. By protecting transgene-derived aberrant RNAs from degradation by the RNA quality control pathway (RQC) in the nucleus, SmD1 presumably promotes cytoplasmic export of transcripts to siRNA bodies, where PTGS occurs (Elvira-Matelot *et al*. 2016). Thus, the involvement of the evolutionarily conserved canonical Sm proteins in splicing-independent posttranscriptional control of mRNA metabolism is emerging as another regulatory component of gene expression. Taken together, these findings suggest that Sm proteins may represent a novel example of RNA-binding proteins that regulate functionally related groups of mRNAs within the cytoplasm. In metabolically active microsporocytes (which enlarge six-fold during diplotene; Kołowerzo-Lubnau *et al*., 2015), the most abundant groups of transcripts are those related to efficient translation, energy production and photosynthesis, which aligns with the presented results. Therefore, it might be assumed that the Sm protein-mRNA interaction within a cell is a ubiquitous process, applying to a vast portion of transcripts.

### The mechanism of Sm protein-mRNA interaction

An open question concerns the mechanism of Sm proteins selectivity toward certain types of mRNAs and mechanism of their interaction. The only Sm proteins-binding elements described to date are the Sm-site present in U snRNAs and yeast telomerase RNA (TLC1 and TER1) (Seto *et al*., 1999; Tang *et al*., 2012), and CAB box found in H/ACA Cajal-body specific RNAs (U85, U87, and U89 scaRNAs) (Darzacq *et al*., 2002; Richard *et al*. 2003) and human telomerase RNA (hTR) (Fu and Collins, 2006). Sm-sites or CAB box domain have not been found in mRNA to date. Furthermore, the specificity of scaRNA binding by Sm proteins is not fully understood. In both *Drosophila* and HeLa cells, Sm proteins co-precipitated only a fraction of the scaRNAs with no apparent structural features that could explain this selectivity (Lu *et al*. 2014). Several studies have demonstrated the participation of spliceosomal U1 snRNPs in splicing-independent posttranscriptional regulation of gene expression, including nascent mRNA protection from premature cleavage and polyadenylation at cryptic sites (Kaida *et al*., 2010; Berg *et al*., 2012), regulation of polyadenylation of retroviral RNA (Ashe *et al*., 1997, 2000; Schrom *et al*., 2013) as well as control of RNA pol II promoter directionality and promotion of sense mRNA transcription (Almada *et al*., 2013; Ntini *et al*., 2013). Recently, it was revealed that, in plants, U1 snRNP regulates miRNA biogenesis (Knop *et al*. 2017; Bhat *et al*. 2019) and mediates the interactions between the microprocessor, spliceosome and polyadenylation machinery (Szweykowska-Kulinska *et al*., 2013; Stepien *et al*., 2016). Moreover, it was shown that, in animal cells, the interaction between Sm proteins and mRNAs is mediated by U1 and presumably other snRNPs. Twelve-nucleotide putative binding sites for the 5’ end of the U1 snRNPs were identified in Sm protein-associated mature mRNAs within the coding sequence far from intron-exon boundaries. It was thus proposed that U1 snRNPs bind mature mRNAs to promote the recruitment of RNA processing factors, thereby affecting mRNA localization, translation and/or turnover (Lu *et al*. 2014). In the case of the larch microsporocytes, the Sm protein-mRNA interaction is apparently not mediated by U snRNPs, since there was no enrichment in U1 snRNA in the RIP-seq analysis. Additionally, the *in situ* localization of splicing elements demonstrated that the U snRNAs and m3G caps did not colocalize within the S-bodies, suggesting a different mechanism of Sm proteins-mRNA interaction in these cells. Studies on *Drosophila* and *C. elegans* have revealed that only some Sm proteins associate with mature mRNAs (Barbee *et al.*, 2002; Gonsalvez *et al.*, 2010; Gonsalvez and Long, 2012). Sm proteins constitute an evolutionarily conserved family and are classified into canonical (spliceosomal) Sm and noncanonical LSm (Sm-like) proteins (Hermann *et al*., 1995; Urlaub *et al*., 2001; Khusial *et al*., 2005; Scofield and Lynch, 2008). In Archaea, LSm proteins participate in tRNA maturation and ribosome synthesis (Mura *et al*., 2003). In Eubacteria, the Hfq homolog of the Sm protein regulates mRNA translation (Schumacher *et al*., 2002). The diversification of the ancestral Sm proteins in eukaryotes enabled their involvement in various processes. The LSm 2-8 ring is found in U6 and U6 atac snRNPs (spliceosomal subunits), as well as in the U8 snoRNPs involved in 5.8S and 28S rRNA processing (Peculis, 1997; Achsel *et al*., 1999). LSm 1-7 form a cytoplasmic complex that promotes mRNA decapping in P-bodies (Bouveret *et al*., 2000, Decker and Parker 2012). Another LSm ring variant, LSm 2-7, binds snR5 snoRNA, which is essential for rRNA pseudouridylation (Fernandez *et al*., 2004). The unique LSm 10 and LSm 11 proteins are part of the U7 snRNP involved in the 3’ end maturation of histone mRNA in animal cells (Schümperli and Pillai, 2004). The *Arabidopsis* genome contains 42 Sm protein and LSM genes (Cao *et al*., 2011), many of which have not yet been characterized. With these findings considered, it could not be precluded that, in larch microsporocytes, the Sm proteins-mRNA association is driven by another noncanonical Sm/LSm ring or is indirectly mediated by other RBPs/RNPs. This hypothesis is supported by an IP-MS analysis that revealed a significant enrichment of the Alba proteins within the cytoplasmic Sm interactome. Alba proteins constitute an evolutionary ancient family of dimeric nucleic acid-binding proteins present in all domains of life (Goyal *et al*. 2016). With their broad functional plasticity, Alba proteins are implicated in genome organization, transcription regulation, RNA metabolism, cell differentiation and stress response. Structurally, plant Alba paralogs display extensive diversity. Their association with target RNA is sequence-independent, and the formation of the Alba-RNA complex facilitates the accessibility of the transcript for RNA processing factors (Goyal *et al*. 2016). It has been shown that the Alba interactome comprises a vast range of different RNAs and RBPs, including mRNA regulators such as PABP, ribosomal subunits and translational factors (Guo *et al*., 2003; Subota *et al*., 2011; Gissot *et al*., 2013). Furthermore, the formation of a particular complex within a cell depends on Alba protein abundance and local concentration, the composition of the dimer (homo or hetero) and posttranslational modifications, including arginine methylation (Lott *et al*., 2013; Ferreira *et al*., 2014). Interestingly, several studies point to a cytoplasmic fraction of the Alba protein within the cell. In *Plasmodium falciparum*, PfAlba 1, 2 and 4 are localized as distinct cytoplasmic foci during proliferation (Chêne *et al*., 2012). Cytoplasmic localization of OsAlba1 was confirmed in rice (*Oryza sativa*), and its level of expression was increased in response to dehydration stress (Verma *et al*., 2014). PAlba 1-3 from *P. berghei* coprecipitate with P granule components PABP and eIF4E (Mair *et al*., 2010). Additionally, the association of TbAlba 1-4 with cytoplasmic mRNAs was demonstrated in *Trypanosoma brucei*, in which it was localized to stress granules upon starvation, indicating their role in spatiotemporal translational regulation (Mani *et al*., 2011; Subota *et al*., 2011). Similarly, the larch microsporocyte cytoplasmic Sm protein interactome was enriched in both the Alba proteins and the ribosomal and translation-related proteins.

### Sm/poly(A) cytoplasmic bodies and the metabolic activity of the microsporocyte

The presented results unambiguously show that the S-bodies accumulate a large pool of mRNAs encoding different proteins. What would be the functional role of this kind of microdomain? The cytoplasm is a highly dynamic cell compartment where the antagonistic processes of mRNA translation and decay are strictly regulated (Rissland, 2017). The spatial separation of the mRNA processing, translational and degradation machinery in distinct microdomains facilitates the efficiency of posttranscriptional regulation and protects the transcripts from uncontrolled translation or degradation, depending on recent needs of the cell (Parker and Sheth, 2007). These studies demonstrate that the S-bodies are not sites of mRNA degradation and therefore are not analogs of P-bodies. This finding is in agreement with previous observations, showing that the mRNAs that accumulate within P-bodies are deadenylated (Sheth and Parker, 2003; Aizer *et al*., 2014), which is not the case with S-bodies.

Our recent studies has shown that the diplotene stage is the most transcriptionally active stage of prophase I (Kołowerzo-Lubnau *et al*., 2015). We comprehensively examined the transcriptional activity, levels and distribution of poly(A) RNA, RNA polymerase II (an enzyme responsible for mRNA synthesis) and snRNAs, which are responsible for mRNA maturation (Smoliński *et al*., 2007, 2011, Smoliński and Kołowerzo 2012; Hyjek *et al*. 2015, Kołowerzo-Lubnau *et al*., 2015). We also examined whether mRNA synthesis and maturation is accompanied by protein synthesis. The periods of intense transcription in larch microsporocytes positively correlated with periods of chromatin decondensation (the diffuse stages), whereas during the chromatin contraction stages, transcription levels decreased. As a result, RNA synthesis is not a continuous process but occurs in transcriptional bursts. The S-bodies were localized after transcriptional bursts, indicating that these microdomains are formed as a result of pulsed, extensive mRNA synthesis in the diffuse stage.

The accumulation of a vast range of mRNAs in spatially separated cytoplasmic bodies is presumably a result of the effective organization of highly expressed transcripts during periods of greatly increased export of these molecules to the cytoplasm. Since S-bodies do not contain markers of degradation, the mRNAs are most likely sequestered to cope with the large number of molecules. This kind of regulation might act as a cellular strategy to control the accessibility of transcripts and their protein products during periods of genome contraction, at which time transcriptional regulation is not possible. This system enables quick expression of proteins essential for normal cell growth and development (preparing the cell for division) and additionally provides the capability to respond to environmental changes and external stimuli. Recent studies have revealed that larch microsporocytes employ an unusual mechanism of nuclear mRNA retention with a putative role for Cajal bodies. It was demonstrated that a significant pool of newly transcribed poly(A) RNAs is temporarily sequestered in the nucleoplasm and CBs before being exported to the cytoplasm (Kołowerzo *et al*., 2009; Smoliński and Kołowerzo, 2012). This functional connection between nuclear mRNA retention, accumulation in Cajal bodies and cytoplasmic localization is further supported by the observation that as many as 13 of the 16 mRNAs investigated as cytoplasmic partners for Sm proteins were also localized to the CBs (see Figure 6, Figure S5).

#### Conclusion

In summary, S-bodies represent newly described intracellular microdomains that participate in mRNA posttranscriptional regulation. Through the temporal storage of mRNAs and their spatial separation from degradation and translation factors, these domains presumably play a role in the coordination of the ongoing, frequently overlapping and competing processes of transcript maturation, translation and degradation. This strict control of mRNA fate is particularly essential in cells such as microsporocytes, in which the chromatin is periodically switched on and off and gene expression occurs in bursts. We propose a model in which the accumulation of significant amounts of different types of mRNA in cytoplasmic bodies provides a mechanism for the temporal retention and storage of highly expressed transcripts until they are needed for further translation or turnover within the cytoplasm, according to the needs of the cell. The results obtained in this work support the hypothesis that evolutionarily conserved Sm proteins have been adapted to perform a plethora of functions related to the posttranscriptional regulation of gene expression in Eukaryota. This multipurpose capacity presumably enabled them to coordinate the interdependent processes of splicing element assembly, mRNA maturation and processing, and mRNA translation regulation and degradation.

## EXPERIMENTAL PROCEDURES

### Plant material

For immunofluorescence/FISH assays, anthers of the European larch (*Larix decidua* Mill.) were collected between November and March at weekly intervals from the same tree in successive meiotic prophase (diplotene) stages to ensure consistent experimental conditions. The anthers were fixed in 4% paraformaldehyde in phosphate-buffered saline (PBS), pH 7.2, for 12 h and squashed to obtain free meiocytes. Meiotic protoplasts were isolated from these cells according to the method of Kołowerzo et al. ^118^. Isolated protoplasts were next subjected to immunostaining, FISH and immuno-FISH assays.

For transmission electron microscopy (TEM) assays, the anthers were fixed in 4% paraformaldehyde and 0.25% glutaraldehyde GA in 0.1 M PIPES, pH 7.2, for 12 h. The samples dedicated for the ultrastructural analysis were additionally fixed with 2% OsO_4_ for 12 h at 4°C. Next, the samples were dehydrated in alcohol and embedded in LR Gold resin (Sigma, St. Louis, Mo., USA). The material was sectioned using a Leica Ultracut UCT ultramicrotome, and the ultrathin sections were placed on nickel-formvar-coated grids. For observations of the ultrastructure, the samples were contrasted with 2.5% uranyl acetate for 30 min and then incubated with 2.5% lead citrate for 15 min.

For immunoprecipitation (IP) and RNA immunoprecipitation (RIP), the larch anthers were flash-frozen in liquid nitrogen and stored at −80°C for further analysis.

### Immunogold labeling of Sm proteins

Ultrathin sections were preincubated with 1% BSA in PBS pH 7.2 for 30 min, followed by incubation with anti-Sm antibody (ANA No. 5, Centers for Disease Control and Prevention, Atlanta, GA 30333, USA) at 1:800 dilution and 4°C overnight. After washing with PBS (5 × 10 min), the samples were incubated with secondary anti-human antibody coupled to 20-nm-diameter colloidal gold particles (BioCell, Cardiff, UK; 1:20 dilution in 1% BSA in PBS, pH 7.2) for 1 h at 37°C, rinsed in PBS and water and then contrasted with 1% phosphotungstic acid (PTA) and 2.5% uranyl acetate. The ultrastructural and immunogold analyses were performed with the use of a JEOL EM 1010 transmission electron microscope.

### Design of the multi-labeling reactions

Prior to the assay, the cells were treated with 0.1% Triton X-100 in PBS for 25 min to induce cell membrane permeabilization. After labeling, the samples were stained for DNA detection with Hoechst (1:2000) and mounted in ProLong Gold Antifade reagent (Life Technologies, Carlsbad, CA, USA). The sequences for all the oligo probes used in this work are summarized in Figure S7.

### Double labeling of the poly(A) RNA/U snRNA/5S rRNA and Sm proteins

The samples were incubated with Cy3-labeled probes: oligo(dT), U1 snRNA, U2 snRNA, U4 snRNA, U5 snRNA and 5S rRNA. The 100 µM stock solutions with the probes were diluted 1:500 in hybridization buffer (30% v/v formamide, 4× SSC, 5× Denhardt’s buffer, 1 mM EDTA, and 50 mM phosphate buffer) and incubated for 4 h (oligo(dT)) or overnight (other probes) at 27°C. After washing with 4× SSC (5 × 1 min), 2× SSC (5 × 1 min) and PBS (1 × 3 min), the samples were treated with the following antibodies for the detection of the Sm proteins: human anti-Sm ANA No. 5 (1:300 in 0.2% acBSA in PBS, pH 7.2), mouse anti-Sm Y12 (Abcam; 1:100 in 0.01% acBSA in PBS, pH 7.2), mouse anti-Sm Y12 (a kind gift from Michael P. Terns, University of Georgia, Athens, GA 30602, USA; 1:500 in 0.01% acBSA in PBS, pH 7.2). The samples were then rinsed with PBS (5 × 3 min) and incubated for 1 h at 37°C with the following secondary antibodies: anti-human Alexa 488 (Thermo Fisher, Waltham, MA, USA; 1:750) or anti-mouse Alexa 488 (Thermo Fisher; 1:1000 in 0.01% acBSA in PBS, pH 7.2).

### Double labeling of the mRNA and Sm proteins

Immunodetection of Sm proteins with ANA No. 5 was performed first to detect the mRNA and Sm proteins according to the procedure described above. The secondary anti-human TRITC antibody (Thermo Fisher; 1:200 in 0.01% acBSA in PBS, pH 7.2) was used in this assay. Next, the samples were incubated overnight at 27°C with DIG-labeled probes that are complementary to the distinct mRNAs (Figure S7) diluted 1:200 in hybridization buffer. After washing with 4× SSC (5 × 1 min), 2× SSC (5 × 1 min), 1× SSC (1 × 10 min) and PBS (1 × 3 min), the slides were incubated for 5 h at 8°C with mouse anti-DIG antibody (Roche, Basel, Basel, Switzerland; 1:200 in 0.05% acBSA in PBS, pH 7.2) and rinsed in PBS. The samples were then incubated for 1 h at 37°C with secondary anti-mouse Alexa 488 at 1:1000 in 0.01% acBSA in PBS, pH 7.2.

### Double localization of the proteins

Samples were incubated overnight at 4°C with the following primary antibodies: mouse anti-m3G (Santa Cruz Biotechnology, Dallas, TX; 1:50), mouse anti-U2B’’ (LifeSpan Bioscience, Seattle, WA, USA; 1:20), rat anti-RNA pol II (Chromotek, Planegg-Martinsried, Germany; elongated form — hyperphosphorylated Ser2 of the CTD; 1:100) and rabbit anti-AGO1 (Agrisera, Vännäs, SWEDEN; 1:200) in 0.01% acBSA in PBS, pH 7.2. After rinsing with PBS, the samples were incubated for 1 h at 37°C with the following secondary antibodies: anti-mouse Alexa 546 (Thermo Fisher), anti-rat Alexa 488 (Thermo Fisher) or anti-rabbit Alexa 488 (Thermo Fisher) 1:500 in 0.01% acBSA in PBS, pH 7.2. Next, the slides were rinsed with PBS, preincubated with 2% BSA in PBS, pH 7.2, for 15 min, and incubated overnight at 4°C with the anti-Sm ANA No. 5 antibody diluted 1:300 in 0.2% acBSA in PBS, pH 7.2. After rinsing with PBS, the samples were incubated for 1 h at 37°C with the appropriate respective anti-human antibody: Alexa 488 at 1:750 or TRITC, at 1:200 in 0.01% acBSA in PBS, pH 7.2.

### Triple labeling of the poly(A) RNA and proteins

For the triple-labeling assays, the oligo(dT) probe was first hybridized according to the method described above (overnight incubation). Next, the samples were incubated overnight at 4°C with the following primary antibodies: mouse anti-DCP2 (Sigma, Saint Louis, MI, USA; 1:20), rabbit anti-LSm4 (Sigma; 1:100) and rabbit anti-RS28 (Sigma; 1:100) in 0.01% acBSA in PBS, pH 7.2. After washing with PBS, the slides were incubated for 1 h at 37°C with anti-mouse Alexa 488 (1:1000) or anti-rabbit Alexa 488 (1:500) in 0.01% acBSA in PBS, pH 7.2. The Sm proteins were labeled as described above with ANA No. 5 primary and anti-human Alexa 633 (Thermo Fisher; 1:200) secondary antibodies.

### Confocal microscopy and image analysis

The images were captured with a Leica TCS SP8 confocal microscope at wavelengths of 405, 488, 543, 561 and 633 nm. The optimized pinhole, long exposure time (400 kHz) and 63X (numerical aperture 1.4) Plan Apochromat DIC H oil immersion lens were used. Images were acquired sequentially in the blue (DAPI), green (Alexa 488), red (TRITC, Cy3, Alexa 546) and/or far red (Alexa 633) channels. The optical sections were collected at 0.5 μm intervals. For the bleed-through analysis and control experiments, LAS AF software was used. For image processing and analysis, ImageJ software was used (Schneider *et al*., 2012).

For the quantitative measurements, each experiment was performed using consistent temperatures, incubation times, and concentrations of probes and antibodies. The images were collected under consistent conditions of acquisition (laser power, emission band, gain, and resolution) to ensure comparable results. Before quantification of the fluorescence intensity, the background was eliminated by adjusting the threshold according to autofluorescence based on the negative control. Between 10 and 30 cells were analyzed for each stage (experimental variant), depending on the experiment. The level of fluorescence was expressed in arbitrary units (as the mean intensity per μm^2^).

The statistical analysis was performed using SigmaPlot 11.0 software. To compare two groups, Student’s t-tests were used. To compare between several groups, statistical significance was determined by the Kruskal-Wallis test. The significance level was set at *p* < 0.05 for all statistical tests.

Colocalization analysis was performed with the ImageJ plugin JACoP (Bolte and Cordelieres, 2006). Between 10 and 25 cells were analyzed for each stage (experimental variant). Only the cytoplasmic area of the cells was subject to all colocalization analyses. The colocalization coefficients were calculated (PCC, Pearson’s correlation coefficient, and M_1_ and M_2_, Manders’ overlap coefficients) for each stage and evaluated for statistical significance with Costes’ randomization test (p < 0.05).

The number and volume of the cytoplasmic bodies per cell were measured with ImageJ plugin 3D Objects Counter (Bolte and Cordelières, 2006). Between 10 and 25 cells for each stage (experimental variant) were analyzed. The statistical significance was determined by one-way ANOVA followed by multiple comparisons using the Holm-Sidak *post hoc* test.

### Immunoprecipitation assays

For proteomic analysis of immunoprecipitates (IP-LC-MS/MS), contents from the cytoplasmic fraction of the cells were immunoprecipitated according to a modified protocol based on (Oliva *et al.* 2016). Briefly, the larch anthers were ground in liquid nitrogen and homogenized in protein extraction buffer at 3 mL per 1 g of plant material. The homogenates were then centrifuged twice at 14 000 × g, followed by centrifugation at 16 000 × g for 30 min at 4°C to obtain native cytosolic protein extract. The supernatant (cytosolic fraction) was precleared with MagnaChiP A/G magnetic beads (Merck) for 1 h at 4°C and incubated with anti-Sm Y12 antibody (a kind gift from Michael P. Terns, University of Georgia, Athens, GA 30602, USA) overnight at 4°C. Next, a fresh set of magnetic beads was added, and the mixture was incubated for 1 h at 4°C. The beads were precipitated by immobilization on a magnetic separator (Thermo Fisher), washed four times with PBST (0.1% Tween 20) and resuspended in water. The experiment was performed in quadruplicate for the IP assay and triplicate for the mock-IP assay (no antibody added). The LC-MS/MS analysis was performed by the Mass Spectrometry Laboratory (IBB PAS, Warsaw) with the use of an Orbitrap mass spectrometer (Thermo Fisher). The proteins were identified with MASCOT software (http://www.matrixscience.com/) and the TAIR10 database. Next, the samples were analyzed with MScan software (http://proteom.ibb.waw.pl/mscan/index.html). The list of Sm protein-immunoprecipitated proteins was corrected with the results from the mock-IP experiments. The protein was considered Sm protein-immunoprecipitated when it was present in at least 3 IP samples and not present in at least 2 control (mock-IP) samples.

For the RNA immunoprecipitation (RIP) from the cytoplasmic fraction, the assays were performed according to a modified protocol from (Sorenson and Bailey-Serres 2015). Briefly, the larch anthers were ground in liquid nitrogen and RNP extraction buffer (200 mM Tris-HCl, pH 9.0; 110 mM potassium acetate; 0.5% (v/v) Triton X-100; 0.1% (v/v) Tween 20; 2.5 mM DTT; complete protease inhibitor (Roche); and 0.04 U/μl RNase inhibitor) was added at 10 mL per 1.5 g of tissue. After thawing on ice, the material was filtered through Miracloth (Merck Millipore, Burlington, MA, USA; ø of 22-25 μm) and centrifuged at 1500 x g for 2 min at 4°C. The supernatant (10 mL per sample) containing the cytoplasmic fraction (Figure S8) of the anther cells was precleared with MagnaChiP A/G magnetic beads for 1 h at 4°C and incubated with anti-Sm antibodies (ANA No. 5 — 15 μl per sample, Y12 (Abcam) — 1 μg per sample, Y12 (Terns) — 10 μl per sample, mock-RIP — no antibody) for 2 h at 4°C. Next, a fresh set of magnetic beads was added and incubated for 1 h at 4°C. The beads were precipitated on a magnetic separator (Thermo Fisher), washed four times with wash buffer (200 mM Tris-HCl, pH 9.0; 110 mM potassium acetate; 0.5% (v/v) Triton X-100; 0.1% (v/v) Tween 20; and 2.5 mM DTT), resuspended in 100 μl of cold wash buffer and subjected to RNA purification and cDNA library preparation.

### cDNA library preparation and sequencing

RNA was purified with a TRIzol:chloroform extraction and incubated with TURBO DNA-free DNase (Thermo Fisher) according to the manufacturer’s protocol. Next, the RNA samples were further purified by phenol:chloroform (Sigma; 25:24) extraction. The RNA pellet was resuspended in 20 μl of RNase-free water.

For qPCR, the RNA was reverse-transcribed with Superscript III (Invitrogen, Carlsbad, CA, USA) enzyme using random hexamers according to the manufacturer’s protocol. The qPCR reaction mix contained 1 µl of template cDNA, 5 µl of 2× concentrated PowerSYBR Green PCR MasterMix (Thermo Fisher), 2 pmoles of each primer and water up to 10 µl. The primers specific for U2 snRNA or 5S rRNA (Supl. 1) were used. The assays were carried out in a 7900HT Fast Real-Time PCR system (Applied Biosystems, Foster City, CA, USA), and the cycling conditions were as follows: 95°C for 10 min; 95°C for 15 s, and 40 cycles of 60°C for 1 min. The relative expression level of the U2 snRNA was calculated via the ΔΔCt method. The results were normalized to the 5S rRNA expression level and compared to the control (RNA isolated from the mock-RIP experiment), for which a value of 1 was assigned.

After qualitative (Bioanalyzer 2100; Agilent) and quantitative (NanoDrop 2000, Qubit; Thermo Fisher) evaluation, the RNA was used for cDNA library preparation with TruSeq Stranded Total RNA with a Ribo-Zero Plant kit (Illumina, San Diego, CA, USA) according to the manufacturer’s protocol. Due to the extremely low RNA amount for the input, 30 cycles of library amplification were performed. The libraries were run on 1.5% agarose gels, and the 250-350 bp fragments were excised, eluted from the gel (GelElute gel extraction kit; Sigma) and quantified via qPCR with a KAPA Library Quantification Kit for Illumina (KAPA Biosystems) according to the manufacturer’s protocol. The libraries were next sequenced on MiSeq (Illumina) with MiSeq Reagent Kit v3 (150 cycles). The reads were filtered in terms of quality via PRINSEQ-lite (Schmieder and Edwards, 2011). The sequence stretches with an average quality <20 over a window of 20 nt were removed, and the reads were cut immediately before the first incidence of a degenerated base. Furthermore, trimmed reads shorter than 36 nt were removed. The reads from all the libraries were pooled and assembled *de novo* with Trinity v.2.8.2 at default settings (Grabherr *et al.*, 2011). The transcript sequences were next analyzed with BLASTX (for putative mRNAs) and BLASTN (for other RNAs) with the use of the NCBI nr and nt databases, respectively. Consensus functions and ontologies were found by LCA (least common ancestor) methodologies implemented in Blast2GO v.4.1 (Götz *et al.*, 2008).

## Supporting information

RIPseq Data

Proteomic data

## ACKNOWLEDGEMENTS

This work was supported by Polish National Science Center NCN grant no. N 303 799640 and grant NCN no 2014/15/N/NZ3/04525.

## REFERENCES

Abdelmohsen, K. Modulation of Gene Expression by RNA Binding Proteins: mRNA Stability and Translation. InTech 123–138. (2012) doi:http://dx.doi.org/10.5772/48485

Achsel, T. et al. A doughnut-shaped heteromer of human Sm-like proteins binds to the 3’-end of U6 snRNA, thereby facilitating U4/U6 duplex formation in vitro. EMBO J. 18, 5789–5802 (1999).

Aizer, A. et al. Quantifying mRNA targeting to P bodies in living human cells reveals a dual role in mRNA decay and storage Accepted manuscript. J. Cell Sci. J. Cell Sci 4443–56 (2014). doi:10.1242/jcs.152975

Almada, A. E., Wu, X., Kriz, A. J., Burge, C. B. and Sharp, A. Promoter directionality is controlled by U1 snRNP and polyadenylation signals. Nature 499, 360–363 (2013).

Anantharaman, V. Koonin, E.V and Aravind, L. (2002) Comparative genomics and evolution of proteins involved in RNA metabolism. Nucleic Acids Res. 30, 1427–1464.

Andrei, M. A. et al. A role for eIF4E and eIF4E-transporter in targeting mRNPs to mammalian processing bodies. RNA 11, 717–727 (2005).

Änkö, M.-L. et al. The RNA-binding landscapes of two SR proteins reveal unique functions and binding to diverse RNA classes. Genome Biol. 13, R17 (2012).

Armstrong, S. J. and Jones, G. H. Meiotic cytology and chromosome behaviour in wild-type Arabidopsis thaliana. J. Exp. Bot. 54, 1–10 (2003).

Ashe, M. P., Furger, A. and Proudfoot, N. J. Stem-loop 1 of the U1 snRNP plays a critical role in the suppression of HIV-1 polyadenylation. RNA 6, 170–7 (2000).

Ashe, M. P., Pearson, L. H. and Proudfoot, N. J. The HIV-1 5’ LTR poly (A) site is inactivated by U1 snRNP interaction with the downstream major splice donor site. EMBO J. 16, 5752–5763 (1997).

Barbee, S. A. and Evans, T. C. The Sm proteins regulate germ cell specification during early C. elegans embryogenesis. Dev. Biol. 291, 132–143 (2006).

Barbee, S. a., Lublin, A. L. and Evans, T. C. (2002) A novel function for the Sm proteins in germ granule localization during C. elegans Embryogenesis. Curr. Biol. 12, 1502–1506.

Baserga, S. J. and Steitz, J.A. The Diverse World of Small Ribonucleoproteins. RNA World, Cold Spring Harb. Lab. Press 359–381 (1993).

Beggs, J. D. Lsm proteins and RNA processing. Biochem. Soc. Trans. 33, 433–438 (2005).

Berg, M. G. et al. U1 snRNP determines mRNA length and regulates isoform expression. Cell 150, 53–64 (2012).

Bhat SS, Bielewicz D, Grzelak N, Gulanicz T, Bodi Z, Szewc L, Bajczyk M, Dolata J, Smolinski DJ, Fray RG, Jarmolowski A, Szweykowska-Kulinska Z (2019). mRNA adenosine methylase (MTA) deposits m6A on pri-miRNAs to modulate miRNA biogenesis in Arabidopsis thaliana. bioRxiv DOI: 10.1101/557900.

Biggiogera, M., Fakan, S., Leser, G., Martin, T. E. and Gordon, J. Immunoelectron microscopical visualization of ribonucleoproteins in the chromatoid body of mouse spermatids. Mol. Reprod. Dev. 26, 150–158 (1990).

Bilinski, S. M., Jaglarz, M. K., Szymanska, B., Etkin, L. D. and Kloc, M. (2004) Sm proteins, the constituents of the spliceosome, are components of nuage and mitochondrial cement in Xenopus oocytes. Exp. Cell Res. 299, 171–178.

Bohn, J. A. et al. Identification of diverse target RNAs that are functionally regulated by human pumilio proteins. Nucleic Acids Res. 46, 362–386 (2018).

Bolte, S. and Cordelières, F. P. A guided tour into subcellular colocalization analysis in light microscopy. J. Microsc. 224, 213–232 (2006).

Borg, M. et al. The R2R3 MYB transcription factor DUO1 activates a male germline-specific regulon essential for sperm cell differentiation in Arabidopsis. Plant Cell 23, 534–49 (2011).

Bouveret, E., Rigaut, G., Shevchenko, A., Wilm, M. and Séraphin, B. A Sm-like protein complex that participates in mRNA degradation. EMBO J. 19, 1661–71 (2000).

Cao, J., Shi, F., Liu, X., Jia, J., Zeng, J., and Huang, G. Genome-wide identification and evolutionary analysis of Arabidopsis sm genes family. J. Biomol. Struct. Dyn. 28, 535–544 (2011).

Cenci, G., Bonaccorsi, S., Pisano, C., Verni, F. and Gatti, M. Chromatin and microtubule organization during premeiotic, meiotic and early postmeiotic stages of Drosophila melanogaster spermatogenesis. J. Cell Sci. 107, 3521–34 (1994).

Cheadle, C. et al. Control of gene expression during T cell activation: alternate regulation of mRNA transcription and mRNA stability. BMC Genomics 6, 75 (2005).

Chêne, A. et al. PfAlbas constitute a new eukaryotic DNA/RNA-binding protein family in malaria parasites. Nucleic Acids Res. 40, 3066–3077 (2012).

Chu, C. Y. and Rana, T. M. Translation repression in human cells by MicroRNA-induced gene silencing requires RCK/p54. PLoS Biol. 4, 1122–1136 (2006).

Chuma, S. et al. (2003) Mouse Tudor Repeat-1 (MTR-1) is a novel component of chromatoid bodies/nuages in male germ cells and forms a complex with snRNPs. Mech. Dev. 120, 979–990.

Colas, I. et al. Observation of Extensive Chromosome Axis Remodeling during the andquot;Diffuse-Phaseandquot; of Meiosis in Large Genome Cereals. Front. Plant Sci. Sci. 8, 1235 (2017).

Comizzoli, P., Pukazhenthi, B. S. and Wildt, D. E. The competence of germinal vesicle oocytes is unrelated to nuclear chromatin configuration and strictly depends on cytoplasmic quantity and quality in the cat model. Hum. Reprod. 26, 2165–2177 (2011).

Darzacq, X. et al. Cajal body-specific small nuclear RNAs: a novel class of 2 ‘-O-methylation and pseudouridylation guide RNAs. Embo J. 21, 2746–2756 (2002).

Decker, C. J. and Parker, R. P-bodies and stress granules: possible roles in the control of translation and mRNA degradation. Cold Spring Harb. Perspect. Biol. 4, (2012).

Elvira-Matelot, E. et al. The nuclear ribonucleoprotein SmD1 interplays with splicing, RNA quality control and post-transcriptional gene silencing in Arabidopsis. Plant Cell 28, 426–438 (2016).

Fernandez, C. F., Pannone, B. K., Xinguo, C. and Fuchs, Gabriele, Wolin, S. L. An Lsm2–Lsm7 Complex in Saccharomyces cerevisiae Associates with the Small Nucleolar RNA snR5. Mol. Biol. Cell 15, 2842–2852 (2004).

Ferreira, T. R. et al. Altered expression of an RBP-associated arginine methyltransferase 7 in Leishmania major affects parasite infection. Mol. Microbiol. 94, 1085–1102 (2014).

Fu, D. and Collins, K. Human telomerase and Cajal body ribonucleoproteins share a unique specificity of Sm protein association. Genes Dev. 20, 531–6 (2006).

Gallo, C. M., Munro, E., Rasoloson, D., Merritt, C. and Seydoux, G. Processing bodies and germ granules are distinct RNA granules that interact in C. elegans embryos. Dev. Biol. 323, 76–87 (2008).

Gama-Carvalho, M., Barbosa-Morais, N. L., Brodsky, A. S., Silver, P. A. and Carmo-Fonseca, M. Genome-wide identification of functionally distinct subsets of cellular mRNAs associated with two nucleocytoplasmic-shuttling mammalian splicing factors. Genome Biol. 7, R113 (2006).

Gerber, A. P., Herschlag, D. and Brown, P. O. Extensive association of functionally and cytotopically related mRNAs with Puf family RNA-binding proteins in yeast. PLoS Biol. 2, 0342–0354 (2004).

Gerber, A. P., Luschnig, S., Krasnow, M. a, Brown, P. O. and Herschlag, D. Genome-wide identification of mRNAs associated with the translational regulator PUMILIO in Drosophila melanogaster. Proc. Natl. Acad. Sci. U. S. A. 103, 4487–92 (2006).

Gibbings, D. J., Ciaudo, C., Erhardt, M. and Voinnet, O. Multivesicular bodies associate with components of miRNA effector complexes and modulate miRNA activity. Nat Cell Biol 11, 1143–1149 (2009).

Gissot, M. et al. Toxoplasma gondii alba proteins are involved in translational control of gene expression. J. Mol. Biol. 425, 1287–1301 (2013).

Goldstrohm, A. C., Hall, T. M. T. and McKenney, K. M. Post-transcriptional Regulatory Functions of Mammalian Pumilio Proteins. Trends Genet. 34, 972–990 (2018).

Gonsalvez, G. B. and Long, R. M. Spatial regulation of translation through RNA localization. F1000 Biol. Rep. 4, 16 (2012).

Gonsalvez, G. B., Rajendra, T. K., Wen, Y., Praveen, K. and Matera, a G. Sm proteins specify germ cell fate by facilitating oskar mRNA localization. Development 137, 2341–2351 (2010).

Gorospe, M. HuR in the mammalian genotoxic response: post-transcriptional multitasking. Cell Cycle 2, 412–414 (2003).

Götz, S. et al. High-throughput functional annotation and data mining with the Blast2GO suite. Nucleic Acids Res. 36, 3420–35 (2008).

Goyal, M., Banerjee, C., Nag, S. and Bandyopadhyay, U. The Alba protein family: Structure and function. Biochim. Biophys. Acta - Proteins Proteomics 1864, 570–583 (2016).

Grabherr, M. G. et al. Full-length transcriptome assembly from RNA-Seq data without a reference genome. Nat. Biotechnol. 29, 644–52 (2011).

Guo, R., Xue, H. and Huang, L. Ssh10b, a conserved thermophilic archaeal protein, binds RNA in vivo. Mol. Microbiol. 50, 1605–1615 (2003).

Halbeisen, R.E. Galgano, A., Scherrer, T. and Gerber, A.P. (2008) Post-transcriptional gene regulation: From genome-wide studies to principles. Cell. Mol. Life Sci. 65, 798–813.

Hermann, H. et al. snRNP Sm proteins share two evolutionarily conserved sequence motifs which are involved in Sm protein-protein interactions. EMBO J. 14, 2076–88 (1995).

Hieronymus, H. and Silver, P. A. A systems view of mRNP biology. Genes Dev. 18, 2845–2860 (2004).

Huang, Y. and Steitz, J. A. Splicing factors SRp20 and 9G8 promote the nucleocytoplasmic export of mRNA. Mol. Cell 7, 899–905 (2001).

Hyjek, M., Wojciechowska, N., Rudzka, M., Kołowerzo-Lubnau, A. and Smoliński, D. J. Spatial regulation of cytoplasmic snRNP assembly at the cellular level. J. Exp. Bot. 66, 7019–7030 (2015).

Jaglarz, M. K., Kloc, M., Jankowska, W., Szymanska, B. and Bilinski, S. M. Nuage morphogenesis becomes more complex: Two translocation pathways and two forms of nuage coexist in Drosophila germline syncytia. Cell Tissue Res. 344, 169–181 (2011).

Kaida, D. et al. U1 snRNP protects pre-mRNAs from premature cleavage and polyadenylation. Nature 468, 664–668 (2010).

Kedersha, N. et al. Stress granules and processing bodies are dynamically linked sites of mRNP remodeling. J. Cell Biol. 169, 871–884 (2005).

Keene, J. D. and Tenenbaum, S. A. Eukaryotic mRNPs may represent posttranscriptional operons. Mol. Cell 9, 1161–1167 (2002).

Keene, J.D. (2007) RNA regulons: coordination of post-transcriptional events. Nat. Rev. Genet. 8, 533–543.

Khusial, P., Plaag, R. and Zieve, G. W. LSm proteins form heptameric rings that bind to RNA via repeating motifs. Trends Biochem. Sci. 30, 522–528 (2005).

Klasterska, I. A new look on the role of the diffuse stage in problems of plant and animal meiosis. 204, 193–203 (1976).

Klasterska, I. The concept of the prophase of meiosis. Hereditas 86, 205–210 (1977).

Knop, K. et al. Active 5 splice sites regulate the biogenesis efficiency of Arabidopsis microRNAs derived from intron-containing genes. Nucleic Acids Res. 45, 2757–2775 (2017).

Kołowerzo, A., Smoliński, D. J. and Bednarska, E. Poly(A) RNA a new component of Cajal bodies. Protoplasma 236, 13–19 (2009).

Kołowerzo-Lubnau, A., Niedojadło, J., Świdziński, M., Bednarska-Kozakiewicz, E. and Smoliński, D. J. Transcriptional activity in diplotene larch microsporocytes, with emphasis on the diffuse stage. PLoS One 10, e0117337 (2015).

Lai, W. S. et al. Evidence that tristetraprolin binds to AU-rich elements and promotes the deadenylation and destabilization of tumor necrosis factor alpha mRNA. Mol. Cell. Biol. 19, 4311–23 (1999).

Lavut, A. and Raveh, D. (2012) Sequestration of highly expressed mrRNAs in cytoplasmic granules, P-bodies, and stress granules enhances cell viability. PLoS Genet. 8, e1002527.

Lee, L., Davies, S. E. and Liu, J. L. (2009) The spinal muscular atrophy protein SMN affects Drosophila germline nuclear organization through the U body-P body pathway. Dev. Biol. 332, 142–155.

Lemaire, R. et al. Stability of a PKCI-1-related mRNA is controlled by the splicing factor ASF / SF2: a novel function for SR proteins. Genes Dev. 16, 594–607 (2002).

Liu, J.-L. and Gall, J. G. (2007) U bodies are cytoplasmic structures that contain uridine-rich small nuclear ribonucleoproteins and associate with P bodies. Proc. Natl. Acad. Sci. U. S. A. 104, 11655–11659.

Liu, Q., Fischer, U., Wang, F. and Dreyfuss, G. The spinal muscular atrophy disease gene product, SMN, and its associated protein SIP1 are in a complex with spliceosomal snRNP proteins. Cell 90, 1013–1021 (1997).

López-Rosas, I. et al. (2012) mRNA Decay Proteins Are Targeted to poly(A)+ RNA and dsRNA-Containing Cytoplasmic Foci That Resemble P-Bodies in Entamoeba histolytica. PLoS One 7, e45966.

Lott, K. et al. Global Proteomic Analysis in Trypanosomes Reveals Unique Proteins and Conserved Cellular Processes Impacted by Arginine Methylation. J Proteomics October 8, 210–225 (2013).

Lü, X., De La Peña, L., Barker, C., Camphausen, K. and Tofilon, P. J. Radiation-induced changes in gene expression involve recruitment of existing messenger RNAs to and away from polysomes. Cancer Res. 66, 1052–1061 (2006).

Lu, Z., Guan, X., Schmidt, C. a and Matera, a G. RIP-seq analysis of eukaryotic Sm proteins identifies three major categories of Sm-containing ribonucleoproteins. Genome Biol. 15, R7 (2014).

Lykke-Andersen, J. and Wagner, E. Recruitment and activation of mRNA decay enzymes by two ARE-mediated decay activation domains in the proteins TTP and BRF-1. Genes Dev. 19, 351–361 (2005).

Mair, G. R. et al. Universal features of post-transcriptional gene regulation are critical for Plasmodium zygote development. PLoS Pathog. 6, (2010).

Mani, J. et al. Alba-domain proteins of trypanosoma brucei are cytoplasmic RNA-Binding proteins that interact with the translation machinery. PLoS One 6, (2011).

Mattaj, I. W. Cap trimethylation of U snRNA is cytoplasmic and dependent on U snRNP protein binding. Cell 46, 905–911 (1986).

Mazan-Mamczarz, K. et al. RNA-binding protein HuR enhances p53 translation in response to ultraviolet light irradiation. Proc. Natl. Acad. Sci. 100, 8354–8359 (2003).

Moser, J.J. and Fritzler, M.J. (2010) Cytoplasmic ribonucleoprotein (RNP) bodies and their relationship to GW/P bodies. Int. J. Biochem. Cell Biol. 42, 828–843.

Moussa, F., Oko, R. and Hermo, L. The immunolocalization of small nuclear ribonucleoprotein particles in testicular cells during the cycle of the seminiferous epithelium of the adult rat. Cell Tissue Res. 278, 363–378 (1994).

Mura, C., Kozhukhovsky, A., Gingery, M., Phillips, M. and Eisenberg, D. The oligomerization and ligand-binding properties of Sm-like archaeal proteins (SmAPs). Protein Sci. 12, 832–847 (2003).

Ntini, E. et al. Polyadenylation site–induced decay of upstream transcripts enforces promoter directionality. Nat. Struct. Mol. Biol. 20, 923–928 (2013).

Ok, S. H. et al. Novel CIPK1-associated proteins in Arabidopsis contain an evolutionarily conserved C-terminal region that mediates nuclear localization. Plant Physiol. 139, 138–50 (2005).

Oliva, M. et al. FIE, a nuclear PRC2 protein, forms cytoplasmic complexes in Arabidopsis thaliana. J. Exp. Bot. 67, 6111–6123 (2016).

Parker, R. and Sheth, U. P Bodies and the Control of mRNA Translation and Degradation. Mol. Cell 25, 635–646 (2007).

Peculis, B. a. The sequence of the 5’ end of the U8 small nucleolar RNA is critical for 5.8S and 28S rRNA maturation. Mol. Cell. Biol. 17, 3702–3713 (1997).

Perea-Resa, C., Hernández-Verdeja, T., López-Cobollo, R., del Mar Castellano, M. and Salinas, J. LSM proteins provide accurate splicing and decay of selected transcripts to ensure normal Arabidopsis development. Plant Cell 24, 4930–4947 (2012).

Raker, V. A., Hartmuth, K., Kastner, B. and Lührmann, R. Spliceosomal U snRNP Core Assembly: Sm Proteins Assemble onto an Sm Site RNA Nonanucleotide in a Specific and Thermodynamically Stable Manner. Mol. Cell. Biol. 19, 6554–6565 (1999).

Richard, P. et al. A common sequence motif determines the Cajal body-specific localization of box H/ACA scaRNAs. EMBO J. 22, 4283–4293 (2003).

Rissland, O. S. The organization and regulation of mRNA-protein complexes. WIREs RNA 8, 1–17 (2017).

Sanford, J. R., Gray, N. K., Beckmann, K. and Cáceres, J. F. A novel role for shuttling SR proteins in mRNA translation A novel role for shuttling SR proteins in mRNA translation. Genes Dev. 18, 755–768 (2004).

Schmieder R and Edwards R. Quality control and preprocessing of metagenomic datasets. Bioinformatics. 27,:863–864 (2011).

Schneider, C. A., Rasband, W. S. and Eliceiri, K. W. NIH Image to ImageJ: 25 years of image analysis. Nat. Methods 9, 671–5 (2012).

Schrom, E.-M. et al. U1snRNP-mediated suppression of polyadenylation in conjunction with the RNA structure controls poly (A) site selection in foamy viruses. Retrovirology 10, 55 (2013).

Schumacher, M. A., Pearson, R. F., Moller, T., Valentin-Hansen, P. and Brennan, R. G. Structures of the pleiotropic translational regulator Hfq and an Hfq-RNA complex: A bacterial Sm-like protein. EMBO J. 21, 3546–3556 (2002).

Schümperli, D. and Pillai, R. S. The special Sm core structure of the U7 snRNP: Far-reaching significance of a small nuclear ribonucleoprotein. Cell. Mol. Life Sci. 61, 2560–2570 (2004).

Scofield, D. G. and Lynch, M. Evolutionary diversification of the Sm family of RNA-associated proteins. Mol. Biol. Evol. 25, 2255–2267 (2008).

Sen, G. L. and Blau, H. M. Argonaute 2/RISC resides in sites of mammalian mRNA decay known as cytoplasmic bodies. Nat. Cell Biol. 7, 633–636 (2005).

Seto, a G., Zaug, a J., Sobel, S. G., Wolin, S. L. and Cech, T. R. Saccharomyces cerevisiae telomerase is an Sm small nuclear ribonucleoprotein particle. Nature 401, 177–80 (1999).

Shaw, D. J., Eggleton, P. and Young, P. J. Joining the dots: Production, processing and targeting of U snRNP to nuclear bodies. Biochim. Biophys. Acta - Mol. Cell Res. 1783, 2137–2144 (2008).

She, W. et al. Chromatin reprogramming during the somatic-to-reproductive cell fate transition in plants. Development 140, 4008–4019 (2013).

Sheehan, M. J. and Pawlowski, W. P. Live imaging of rapid chromosome movements in meiotic prophase I in maize. Proc. Natl. Acad. Sci. U. S. A. 106, 20989–94 (2009).

Sheth, U. and Parker, R. (2003) Decapping and Decay of Messenger RNA Occur in Cytoplasmic Processing Bodies. Science. 300, 805–808.

Sieburth, L. E. and Vincent, J. N. Beyond transcription factors: roles of mRNA decay in regulating gene expression in plants. F1000Research 7, 1940 (2018).

Smoliński, D. J. and Kołowerzo, A. mRNA accumulation in the Cajal bodies of the diplotene larch microsporocyte. Chromosoma 121, 37–48 (2012).

Smoliński, D. J., Niedojadło, J., Noble, A. and Górska-Brylass, A. Additional nucleoli and NOR activity during meiotic prophase I in larch (Larix decidua Mill.). Protoplasma 232, 109–120 (2007).

Smoliński, D. J., Wróbel, B., Noble, A., Zienkiewicz, A. and Górska-Brylass, A. Periodic expression of Sm proteins parallels formation of nuclear cajal bodies and cytoplasmic snRNP-rich bodies. Histochem. Cell Biol. 136, 527–541 (2011).

Sorenson, R. and Bailey-Serres, J. Rapid immunopurification of ribonucleoprotein complexes of plants. Methods Mol Biol. 1284, 209–219 (2015).

Stepien, A. et al. Posttranscriptional coordination of splicing and miRNA biogenesis in plants. WIREs RNA doi:10.10, (2016).

Subota, I. et al. ALBA proteins are stage regulated during trypanosome development in the tsetse fly and participate in differentiation. Mol. Biol. Cell 22, 4205–4219 (2011).

Szweykowska-Kulinska, Z., Jarmolowski, A. and Vazquez, F. The crosstalk between plant microRNA biogenesis factors and the spliceosome. Plant Signal. Behav. 8, e26955 (2013).

Tam, P. pc et al. The Puf family of RNA-binding proteins in plants: phylogeny, structural modeling, activity and subcellular localization. BMC Plant Biol. 10, 44 (2010).

Tang, W., Kannan, R., Blanchette, M. and Baumann, P. Telomerase RNA biogenesis involves sequential binding by Sm and Lsm complexes. Nature 484, 260–264 (2012).

Ule, J. et al. CLIP Identifies Nova-Regulated RNA Networks in the Brain. Science 80. 302, 1212–1215 (2003).

Unterholzner, L. and Izaurralde, E. SMG7 acts as a molecular link between mRNA surveillance and mRNA decay. Mol. Cell 16, 587–596 (2004).

Urlaub, H., Hartmuth, K., Kostka, S., Grelle, G. and Lührmann, R. A general approach for identification of RNA-protein cross-linking sites within native human spliceosomal small nuclear ribonucleoproteins (snRNPs): Analysis of RNA-protein contacts in native U1 and U4/U6.U5 snRNPs. J. Biol. Chem. 275, 41458–41468 (2000).

Urlaub, H., Raker, V. A., Kostka, S. and Lührmann, R. Sm protein-Sm site RNA interactions within the inner ring of the spliceosomal snRNP core structure. EMBO J. 20, 187–196 (2001).

Verma, J. K. et al. OsAlba1, a dehydration-responsive nuclear protein of rice (Oryza sativa L. ssp. indica), participates in stress adaptation. Phytochemistry 100, 16–25 (2014).

Wilczynska, a, Aigueperse, C., Kress, M., Dautry, F. and Weil, D. The translational regulator CPEB1 provides a link between dcp1 bodies and stress granules. J. Cell Sci. 118, 981–992 (2005).

Xu, J. and Chua, N.-H. Arabidopsis decapping 5 is required for mRNA decapping, P-body formation, and translational repression during postembryonic development. Plant Cell 21, 3270–3279 (2009).

Yong, J., Kasim, M., Bachorik, J. L., Wan, L. and Dreyfuss, G. Gemin5 Delivers snRNA Precursors to the SMN Complex for snRNP Biogenesis. Mol. Cell 38, 551–562 (2010).

Zhang, S. G. et al. Development of male gametophyte of Larix leptolepis Gord. with emphasis on diffuse stage of meiosis. Plant Cell Rep. 27, 1687–1696 (2008).

Zieve, G. W., Sauterer, R. A. and Feeney, R. J. Newly synthesized small nuclear RNAs appear transiently in the cytoplasm. J. Mol. Biol. 199, 259–267 (1988).

