## Supplementary Figures 1-8 for "Core spliceosomal Sm proteins as constituents of cytoplasmic mRNPs in plants"

**Figure S1 Localization of the Sm proteins and poly(A) RNA on semithin sections of anther *Larix decidua* Mill.**

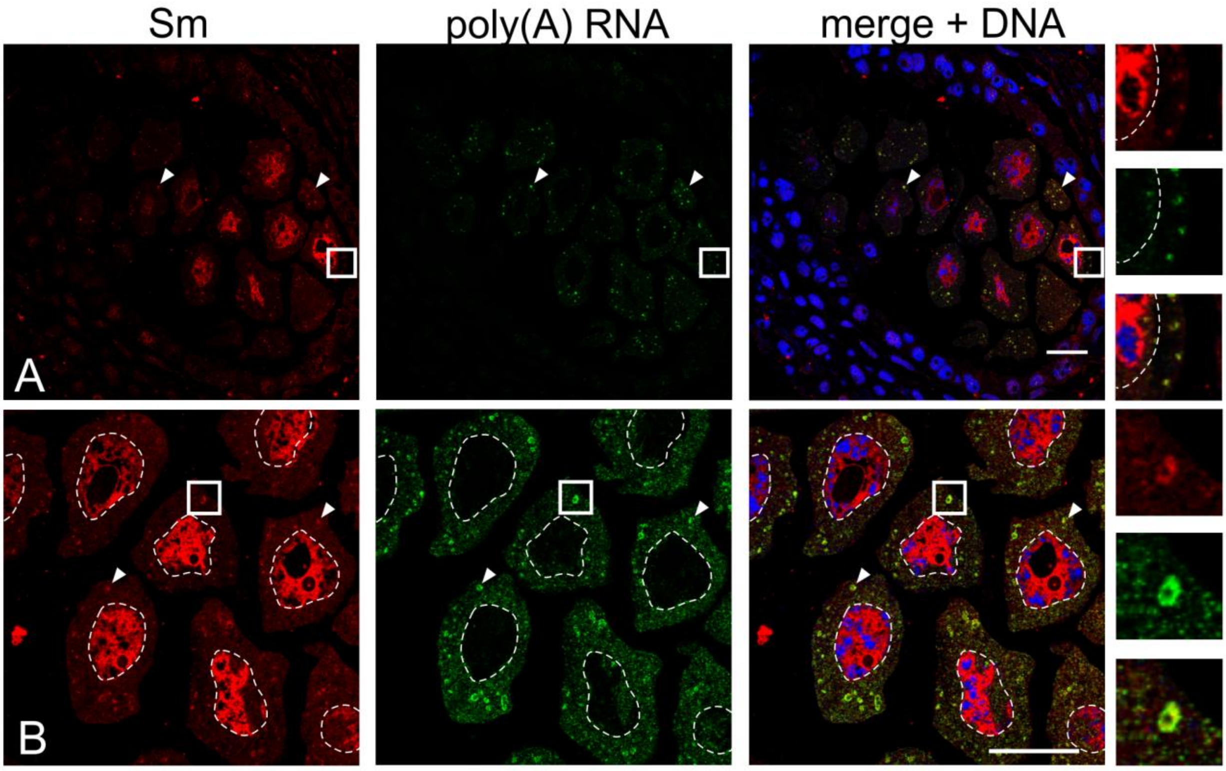


In the cytoplasm, visible colocalizing fluorescent foci (arrowheads). Bar - 25 µm.

**Figure S2 Analysis of Sm protein distribution during colocalization with selected splicing U snRNAs.**

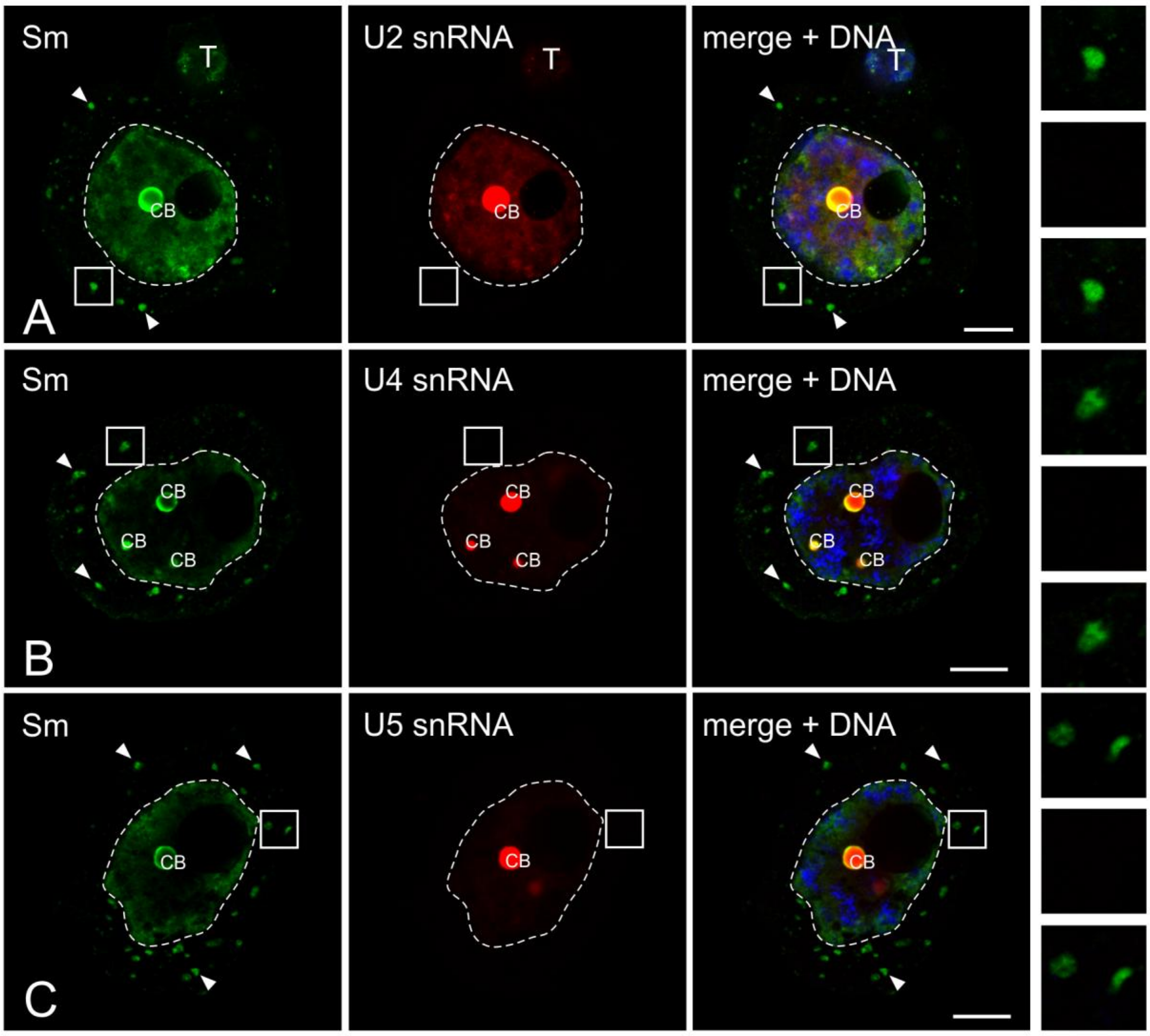


**A,** U2 snRNA; **B**, U4 snRNA; **C**, U5 snRNA. There was no accumulation of U snRNAs in the S-bodies (arrowheads). The right panel represents the magnification of the cytoplasm, which is marked with a square. CB - Cajal body. Bar - 10 μm.

**Figure S3** Western blot analysis of protein extracts obtained after immunoprecipitation using different types of anti-Sm antibodies (Y12, ANA No.5, Y12 (Abcam)). nz - fraction not bound by the antibody. K- - control without antibody. input - total cytoplasmic extract of Larix decidua anther cells. Detection of Sm proteins on the membrane was performed using ANA No. 5.

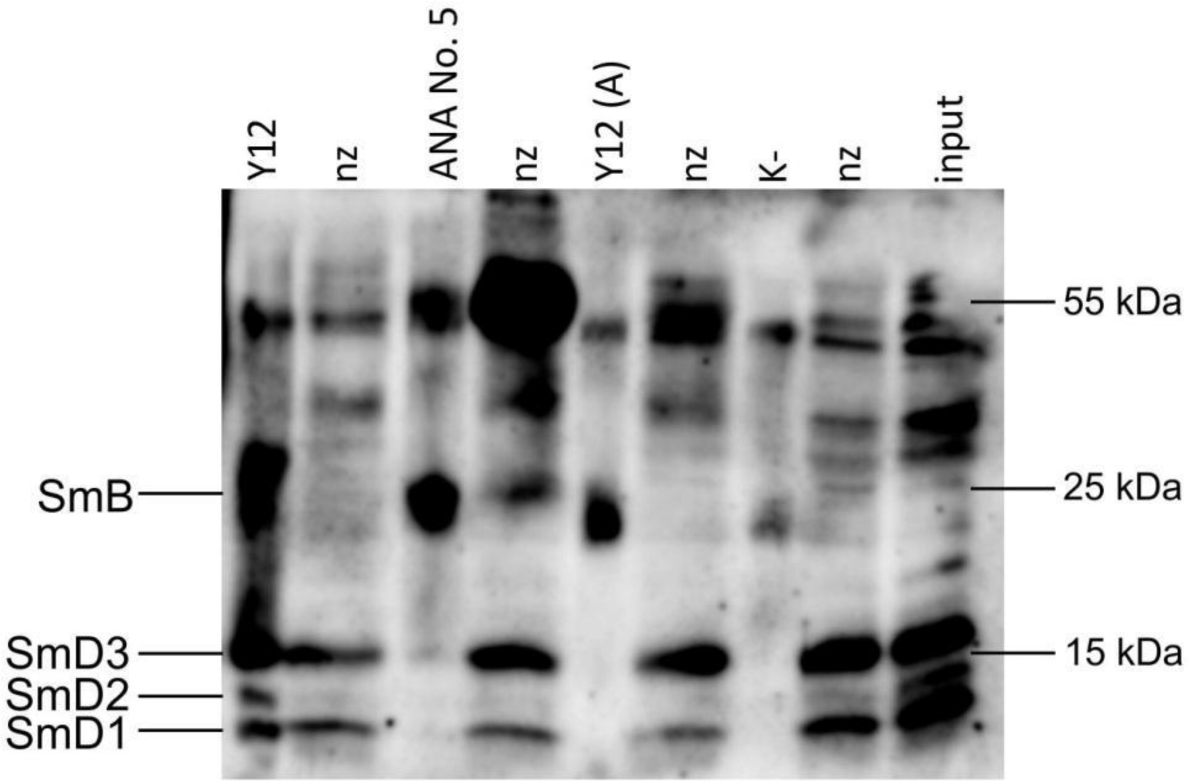

**Fig S4 Sm proteins are absent from control RNA immunoprecipitations and present in Y12 precipitations**

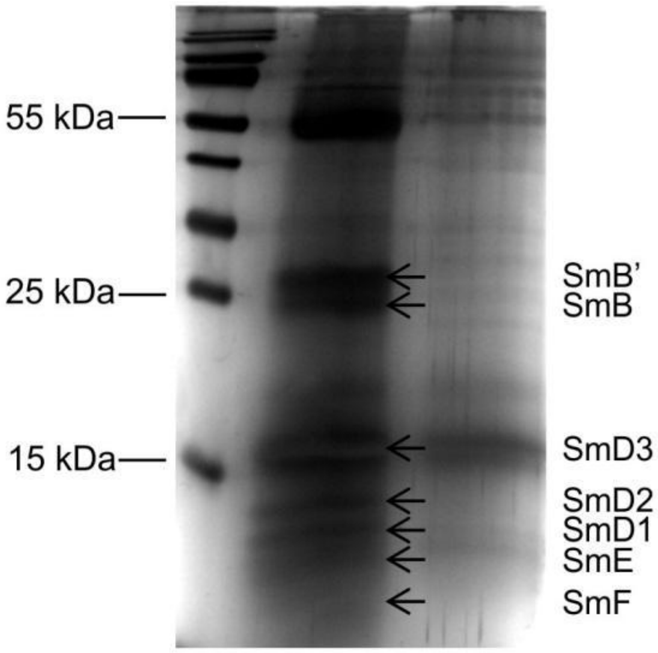


Identification of Sm Larix decidua proteins by IP-SDS-PAGE-MS. Marker (left panel), IP (central panel)- sample after immunoprecipitation using Y12 antibodies, control (right panel) - sample after "immunoprecipitation" without antibody. Individual Sm proteins were identified on the basis of mass spectrometry (MS) analysis cut from the bands.

**Figure S5**

**Localization of mRNA precipitating with Sm proteins *in situ*.**

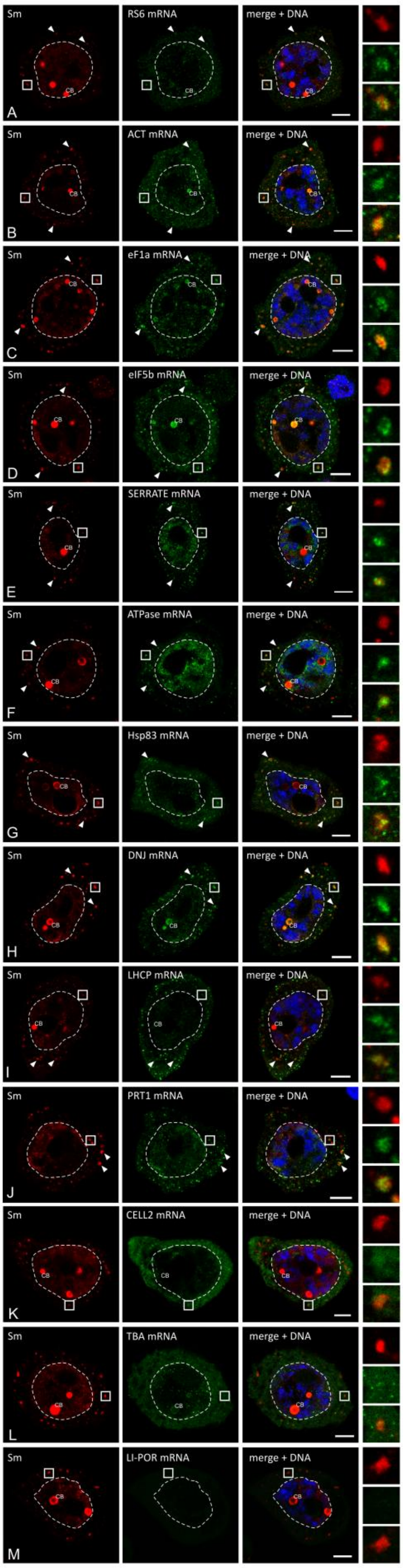


The results presented represent the stages of the IV - VI poly(A) RNA cycle. Numerous clusters of mRNA that colocalized with Sm proteins (caves) are visible in the cytoplasm. The right panel represents the magnification of the cytoplasm, which is marked with a square. RS6 - small ribosome subunit protein S6; RL 37a - large ribosome subunit protein 37a; eF1a - translation factor 1a; eIF5b - translation factor 5b; ATPase - ATPase; KIN12b - kinesin 12b-like protein; LHCP - chlorophyll binding protein ab; NAD7 - NADH 7 dehydrogenase subunit, PRT1 - ubiquitin-dependent ligand PRT1; ACT - actin; Hsp83 - heat shock protein 83; DNJ - protein with DNaJ domain; and CB - Cajal body. Bar 10 µm.

**Figure S6**

**Localization of the Sm mRNAs and Sm proteins *in situ*.**

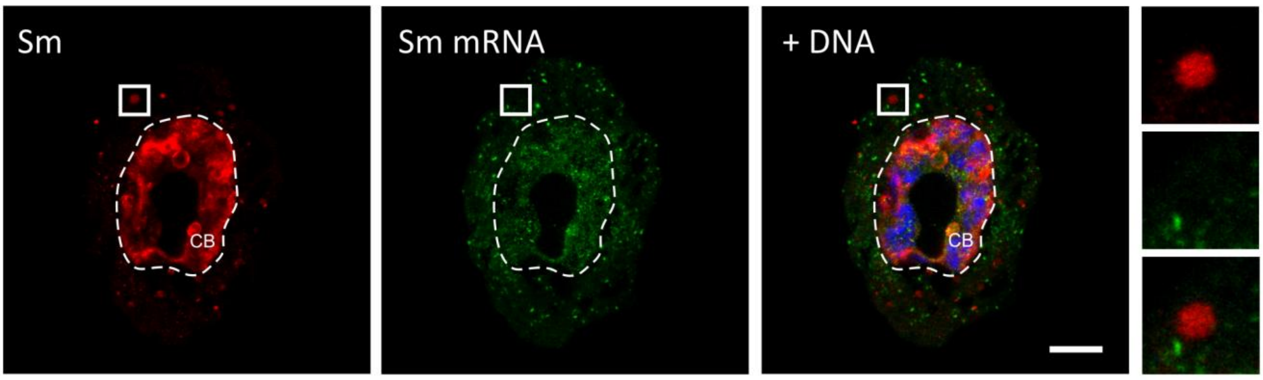


The results presented represent the stages of the IV - VI poly(A) RNA cycle. Numerous clusters of fluorescence showing no colocalization with the Sm proteins, can be seen in the cytoplasm. The right panel represents the magnification of the cytoplasm, which is marked with a square. CB - Cajal body. Bar 10 µm.

**Figure S7 Real-time RT PCR primers and probe sequences used for *in situ* hybridization.**

| Name | Sequence (5' – 3') | Labeling |
|---|---|---|
| **PCR** | | |
| F_U2 snRNA | ATACCTTTCTCGGCCTTTTGGC | - |
| R_U2 snRNA | CAGGCTNGTGCTNTAGTGCAAC | - |
| F_5S rRNA | GTGCGATCATACCAGCGTTA | - |
| R_5S rRNA | CCCATCCCAGCACTACTCTA | - |
| ***in situ* hybridization** | | |
| poly(A) | $T(T)_{29}$ | Cy3 |
| U1 snRNA | TCAGTGGGAGATGCTAGGTAGAC | Cy3 |
| U2 snRNA | ATATTAAACTGATAAGAACAGATACTACACTTG | Cy3 |
| U4 snRNA | GGAAATAGTTTTCAACCAGCAATAGAC | Cy3 |
| U5 snRNA | TATTCTTTAGTAAAAGGCGAAAGAATAGTT | Cy3 |
| 5S rRNA | GAGGTCACCCATCCTAGTACTACTCTC | Cy3 |
| RS6 mRNA | TTCCTTTGCAGAGTCAATGGTGTCACAAGCC | DIG |
| RL 37a mRNA | TGGCCATTACTCTTCTTCACCTTCTTCAGCTT | DIG |
| eF1a mRNA | TGGTGACCTTTGCACCAGTGGGATCCTTCT | DIG |
| eIF5b mRNA | TCTTCACACCAAGTTCATCTGCAAGCTCACGAG | DIG |
| SERRATE mRNA | GTTCCATAGTAATCCAATCCATGAACTCGCCA | DIG |
| ATPase mRNA | ACCATGTAGTGTCCTTTGAGCATGTGCCCATT | DIG |
| KIN12B mRNA | TTCATTGACAATTGCCTTATTCTGTATCGCCTT | DIG |
| LHCP mRNA | AACGCAGCCAAGCGCCCCCAACATGGCCCATCT | DIG |
| NAD7 mRNA | TCCATGGATAGTTTCATTTGACATCGTGATGG | DIG |
| PRT1 mRNA | TGTTGAGAGAAGCCTAATCAGACCCGTGTA | DIG |
| ACT mRNA | TCTTCTGGAGCAACTCGAAGCTCATTGTAGA | DIG |
| Hsp83 mRNA | ACAGTCATAACACCACCTGCTGTCTCGAGAC | DIG |
| DNJ mRNA | TCTTTAAGGGCATCCTCTCCATACTGGTCA | DIG |
| PEP mRNA | TGTAACCCGTGCATAAGGTCCACTCGAACGTGC | DIG |
| PER40 mRNA | TGAATGTTGCACAACGTGCCTTTCCTATTGTA | DIG |
| ADPRF mRNA | TCATCCCTTGCTTCTACAACACGGTCTCTGT | DIG |
| CLATH mRNA | TGCAAGATTCACAGCAAGCTCTAGATTATTCAGC | DIG |
| SDH mRNA | ACTTGTTCGTCTCTTCATAGACCTCCCCACCAAT | DIG |
| SNRP27 mRNA | TCGGAATCCCCAATTTCTTCATCATCTC | DIG |
| ASIL2 mRNA | TCCAGATCCATCATCTGCTGATGCTTCACCA | DIG |
| PABP4 mRNA | ACTCCATAAAGCCATAGCCCTCTGACTGTCCA | DIG |
| CALX mRNA | TCGTAGCTTCAGGATCGTCAATCTCCTCGG | DIG |
| SSL3 mRNA | ATCGGAGTTCTACAACAGTCTATTGCAACC | DIG |
| PLY mRNA | TCGCAACGCCAGCAATCATCAATGGGGTTACCAGT | DIG |
| Sm mRNA | AGGAGTTGACATAAGTATGAATACACATCT | DIG |
| | CTATTCACTTTGGATTTCTAAGAACAATAAT | |
| | GTATATAATTTCCTCTTCAAAAACTACTTTTCTCT | |
| | AATACAAGGTTCATGTACTCATCAAATC | |
| CELL2 mRNA | TTCCGATTGCAGAGAGTCACTGTAAGCGCCT | DIG |
| TBA mRNA | ACTGTTGGTGATCTCGGCTACCGAGAGCT | DIG |
| LI-POR mRNA | AGGTGTCAATATGATCAGATTGGGACCTTCCTC | DIG |

**Figure S8  Validation of the *Larix decidua* anther cell fractionation method.**

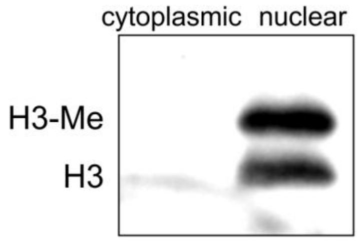


Identification of histone H3 in the cytoplasmic and nuclear fractions by Western blotting. No signal was detected in the cytoplasmic fraction.
