## Supplementary material for "Core spliceosomal Sm proteins as constituents of cytoplasmic mRNPs in plants": Proteomic data

**Supplementary Table 1 Identification of the proteins that formforming a complex with Sm proteins in the cytoplasm determined by IP-LC-MS / MS.**

| No. | locus | protein | score (# of unique peptides) |
|---|---|---|---|
| 1 | AT5G60160.1 | Zn-dependent exopeptidases superfamily protein | 2076.0 (4) |
| 2 | AT4G02840.1,AT4G02840.2 | Small nuclear ribonucleoprotein family protein D1b (SmD1b) | 1692.0 (5) |
| 3 | AT3G07590.1,AT3G07590.2 | Small nuclear ribonucleoprotein family protein D1a (SmD1a) | 1551.0 (3) |
| 4 | AT1G03330.1 | Small nuclear ribonucleoprotein family protein (LSM2) | 1153.0 (1) |
| 5 | AT4G37930.1 | Serine transhydroxymethyltransferase 1 (SHM1, STM, SHMT1) | 1104.0 (10) |
| 6 | AT5G26780.1,AT5G26780.2,AT5G26780.3 | Serine hydroxymethyltransferase 2 (SHM2) | 1104.0 (7) |
| 7 | AT1G76300.1 | snRNP core protein (SmD3) | 1011.0 (3) |
| 8 | AT1G20580.1 | Small nuclear ribonucleoprotein family protein D3b (SmD3b) | 1011.0 (2) |
| 9 | AT1G29250.1,AT2G34160.1 | Alba DNA/RNA-binding protein | 650.0 (4) |
| 10 | AT3G04620.1 | Alba DNA/RNA-binding protein | 650.0 (3) |
| 11 | AT3G13060.1,AT3G13060.2 | Evolutionarily conserved C-terminal region 5 (ECT5) | 634.0 (3) |
| 12 | AT5G27720.1 | Sm-like protein (LSM4) | 509.0 (3) |
| 13 | AT5G20320.1,AT5G20320.2 | Dicer-like protein 4 (DCL4) | 496.0 (7) |
| 14 | AT2G36250.1,AT2G36250.2 | Tubulin/FtsZ family protein (FTSZ2-1) | 439.0 (5) |
| 15 | AT5G04710.1 | Zn-dependent exopeptidases superfamily protein | 438.0 (5) |
| 16 | AT3G52750.1 | Tubulin/FtsZ family protein (FTSZ2-2) | 326.0 (3) |
| 17 | AT2G45290.1 | Transketolase 2 (TKL2) | 290.0 (4) |
| 18 | AT1G26230.1 | Chaperonin 60 subunit beta 4 (CPN-60 beta 4) | 273.0 (2) |
| 19 | AT1G26230.2 | Chaperonin 60 subunit beta 4 (CPN-60 beta 4) | 273.0 (3) |
| 20 | AT4G00620.1 | EMBRYO DEFECTIVE 3127 (EMB3127) | 272.0 (2) |
| 21 | AT4G00600.1 | EMBRYO DEFECTIVE 3127 (EMB3127) | 272.0 (2) |
| 22 | AT4G20440.1,AT4G20440.2,AT4G20440.3, AT4G20440.4,AT5G44500.1,AT5G44500.2 | Small nuclear ribonucleoprotein family protein B (SmB) | 259.0 (3) |
| 23 | AT3G07920.1 | Translation initiation factor IF2/IF5 | 258.0 (2) |
| 24 | AT2G04842.1 | EMBRYO DEFECTIVE 2761 (EMB2761) | 248.0 (4) |
| 25 | AT3G43190.1 | SUCROSE SYNTHASE 4 (SUS4) | 241.0 (5) |
| 26 | AT5G20830.1,AT5G20830.2 | SUCROSE SYNTHASE 1 (SUS1) | 241.0 (7) |
| 27 | AT5G28000.1 | Polyketide cyclase/dehydrase and lipid transport superfamily protein | 230.0 (1) |
| 28 | AT5G49190.1 | SUCROSE SYNTHASE 2 (SUS2) | 224.0 (2) |
| 29 | AT4G02280.1 | SUCROSE SYNTHASE 3 (SUS3) | 224.0 (5) |
| 30 | AT1G79270.1,AT5G61020.1,AT5G61020.2 | Evolutionarily conserved C-terminal region 8 (ECT8) | 219.0 (3) |
| 31 | AT4G19045.1,AT5G45550.1 | MOB kinase activator-like 1 (MOB1-LIKE) | 201.0 (2) |
| 32 | AT1G76860.1 | Small nuclear ribonucleoprotein family protein (LSM3B) | 200.0 (3) |
| 33 | AT1G60110.1 | Mannose-binding lectin superfamily protein | 193.0 (2) |
| 34 | AT1G60130.1 | Mannose-binding lectin superfamily protein | 193.0 (2) |
| 35 | AT3G53510.1 | ABC-2 type transporter family protein (ABCG20) | 184.0 (4) |
| 36 | AT1G78370.1 | glutathione S-transferase TAU 20 (GSTU20) | 176.0 (2) |
| 37 | AT1G21190.1 | Small nuclear ribonucleoprotein family protein (LSM3A) | 175.0 (3) |
| 38 | AT2G18740.1,AT2G18740.2,AT4G30330.1 | Small nuclear ribonucleoprotein family protein | 172.0 (2) |
| 39 | AT4G34200.1 | D-3-phosphoglycerate dehydrogenase 1 (PGDH1) | 163.0 (6) |
| 40 | AT5G22440.1,AT5G22440.2 | Ribosomal protein L1p/L10e family (RL10e) | 160.0 (4) |
| 41 | AT5G26860.1 | lon protease 1 (LON1) | 153.0 (4) |
| 42 | AT1G76950.1 | Regulator of chromosome condensation (RCC1) family (PRAF1) | 151.0 (4) |
| 43 | AT2G39730.1,AT2G39730.2,AT2G39730.3 | Rubisco activase (RCA) | 145.0 (4) |
| 44 | AT2G16440.1 | Minichromosome maintenance (MCM2/3/5) family protein (MCM4) | 140.0 (4) |
| 45 | AT3G54640.1 | Tryptophan synthase alpha chain (TSA1, TRP3) | 123.0 (3) |
| 46 | AT2G35120.1 | Single hybrid motif superfamily protein | 118.0 (2) |
| 47 | AT4G04730.1 | Unknown protein | 118.0 (4) |
| 48 | AT2G17130.1,AT2G17130.2,AT4G35260.1 | Isocitrate dehydrogenase subunit (IDH) | 111.0 (3) |
| 49 | AT5G64140.1 | Ribosomal protein S28 (RPS28) | 110.0 (1) |
| 50 | AT3G45450.1,AT3G48870.1,AT3G48870.2 | ClpC-like protein | 108.0 (2) |
| 51 | AT1G75600.1 | Histone superfamily protein (HTR14) | 107.0 (3) |
| 52 | AT2G23420.1,AT4G36940.1 | Nicotinate phosphoribosyltransferase (NAPRT) | 107.0 (5) |
| 53 | AT5G46560.1 | Inner nuclear membrane protein (MAN1) | 107.0 (3) |
| 54 | AT1G27280.1 | Paired amphipathic helix (PAH2) superfamily protein | 107.0 (2) |
| 55 | AT3G24080.1,AT3G24080.2 | KRR1 family protein | 104.0 (1) |
| 56 | AT4G27070.1 | Tryptophan synthase beta-subunit 2 (TSB2) | 102.0 (3) |
| 57 | AT1G64440.1 | UDP-GLUCOSE 4-EPIMERASE (RHD1, REB1, UGE4) | 100.0 (3) |
| 58 | AT3G22960.1 | Pyruvate kinase family protein (PKP1, PKP-ALPHA) | 95.0 (7) |
| 59 | AT3G60750.1,AT3G60750.2 | Transketolase (TKL1) | 92.0 (4) |
| 60 | AT1G20950.1,AT1G76550.1 | Phosphofructokinase family protein | 90.0 (3) |
| 61 | AT5G49030.1,AT5G49030.2,AT5G49030.3 | tRNA synthetase class I (I, L, M and V) family protein (OVA2) | 87.0 (10) |
| 62 | AT3G03120.1 | ADP-ribosylation factor B1C (ARFB1C) | 84.0 (2) |
| 63 | AT5G17060.1 | ADP-ribosylation factor B1B (ARFB1B) | 84.0 (3) |
| 64 | AT5G60930.1 | P-loop containing nucleoside triphosphate hydrolases superfamily protein | 83.0 (7) |
| 65 | AT3G55510.1 | Noc2p family (RBL) | 81.0 (1) |
| 66 | AT3G25820.2 | Terpene synthase-like sequence-1,8-cineole (TPS-CIN) | 79.0 (1) |
| 67 | AT1G12920.1,AT5G47880.1,AT5G47880.2 | Eukaryotic release factor 1 (ERF1) | 79.0 (4) |
| 68 | AT4G08320.1,AT4G08320.2 | Tetratricopeptide repeat (TPR)-like superfamily protein (TPR8) | 75.0 (1) |
| 69 | AT2G10260.1,AT2G10260.2 | unknown protein, similar to Ulp1 protease family protein (TAIR:AT5G45570.1) | 74.0 (3) |
| 70 | AT5G30510.1 | Ribosomal protein S1 (RPS1) | 70.0 (3) |
| 71 | AT1G50770.1 | Aminotransferase-like, plant mobile domain family protein | 68.0 (4) |
| 72 | AT1G01950.1,AT1G01950.2,AT1G01950.3 | Armadillo repeat kinesin 2 (ARK2) | 63.0 (2) |
| 73 | AT1G59990.1 | EMBRYO DEFECTIVE 3108, DEAD-box ATP-dependent RNA helicase 22 | 61.0 (3) |
| 74 | AT4G40020.1 | Myosin heavy chain-related protein | 61.0 (3) |
| 75 | AT3G50240.1 | Kinesin-related protein (KICP-02) | 61.0 (5) |
| 76 | AT3G44716.1,AT3G44716.2 | unknown protein | 61.0 (1) |
| 77 | AT2G17930.1 | Phosphatidylinositol 3- and 4-kinase family protein with FAT domain | 60.0 (8) |
| 78 | AT1G30440.1 | Phototropic-responsive NPH3 family protein | 60.0 (3) |
| 79 | AT5G22450.1 | unknown protein | 57.0 (6) |
| 80 | AT4G32820.1,AT4G32820.2 | CALCINEURIN BINDING PROTEIN 1 (CABIN1) | 57.0 (5) |
| 81 | AT4G36760.1 | Aminopeptidase P1 (APP1) | 54.0 (3) |
| 82 | AT1G15690.1,AT1G15690.2 | Inorganic H pyrophosphatase family protein (AVP1, AVP-3) | 54.0 (1) |
| 83 | AT5G61950.1 | Ubiquitin carboxyl-terminal hydrolase-related protein | 53.0 (7) |
| 84 | AT1G06590.1 | unknown protein | 51.0 (3) |
| 85 | ATMG00090.1 | ribosomal protein S3 (RS3) | 47.0 (2) |
| 86 | AT5G58530.1 | Glutaredoxin family protein | 47.0 (1) |
| 87 | AT1G11990.1 | O-fucosyltransferase family protein | 47.0 (4) |
| 88 | AT3G06720.1,AT3G06720.2,AT4G16143.1, AT4G16143.2 | importin alpha isoform (IMPA) | 46.0 (1) |
| 89 | AT2G36720.1 | Acyl-CoA N-acyltransferase with RING/FYVE/PHD-type zinc finger domain | 46.0 (3) |
| 90 | AT3G48190.1 | PCD IN MALE GAMETOGENESIS 1 (PIG1) | 46.0 (15) |
| 91 | AT3G44050.1 | P-loop containing nucleoside triphosphate hydrolases superfamily protein | 45.0 (7) |
| 92 | AT1G12470.1 | zinc ion binding protein | 45.0 (2) |
| 93 | AT5G08390.1 | Transducin/WD40 repeat-like superfamily protein | 45.0 (5) |
| 94 | AT1G48300.1 | Diacylglycerol acyltrasferase 3 (DGAT3) | 45.0 (2) |
| 95 | AT5G23340.1 | RNI-like superfamily protein | 43.0 (1) |
| 96 | AT3G01310.1,AT3G01310.2 | Arabidopsis homolog protein of yeast VIP1 1 | 43.0 (4) |
| 97 | AT5G15070.1,AT5G15070.2 | Arabidopsis homolog protein of yeast VIP1 2 | 43.0 (3) |
| 98 | AT3G53940.1 | Mitochondrial substrate carrier family protein | 42.0 (2) |
| 99 | AT4G35350.1 | Xylem cysteine peptidase 1 (XCP1) | 41.0 (2) |
| 100 | AT5G35604.1 | similar to: myosin heavy chain-related (TAIR:AT5G32590.1) | 39.0 (4) |
| 101 | AT1G67120.1 | Midasin | 38.0 (18) |
| 102 | AT3G10220.1 | EMBRYO DEFECTIVE 2804, tubulin folding cofactor B | 35.0 (2) |
| 103 | AT3G04960.1,AT3G04960.2,AT3G04960.3, AT3G04960.4 | Domain of unknown function (DUF3444) | 35.0 (2) |
| 104 | AT4G15360.1 | cytochrome P450, family 705, subfamily A, polypeptide 3 (CYP705A3) | 34.0 (8) |
| 105 | AT3G54050.1,AT3G54050.2 | High cyclic electron flow 1 (HCEF1) | 34.0 (1) |
| 106 | AT1G54460.1 | TPX2 (targeting protein for Xklp2) protein family | 33.0 (1) |
| 107 | AT2G36460.1,AT2G36460.2 | Aldolase superfamily protein | 33.0 (1) |
| 108 | AT4G29640.1 | Cytidine/deoxycytidylate deaminase family protein | 32.0 (4) |
| 109 | AT1G48240.1,AT3G17440.1,AT3G17440.2 | novel plant snare 12 (NPSN12) | 31.0 (2) |
| 110 | AT5G05320.1 | FAD/NAD(P)-binding oxidoreductase family protein | 30.0 (1) |
| 111 | AT4G28630.1 | ABC transporter of the mitochondrion 1 (ATM1) | 29.0 (1) |
| 112 | AT1G16400.1 | cytochrome P450, family 79, subfamily F, polypeptide 2 (CYP79F2) | 29.0 (2) |
| 113 | AT5G61150.1,AT5G61150.2 | VERNALIZATION INDEPENDENCE 4 (VIP4) | 28.0 (4) |
| 114 | AT2G38720.1 | Microtubule-associated protein 65-5 (MAP65-5) | 28.0 (1) |
| 115 | AT1G67950.2 | RNA-binding (RRM/RBD/RNP motifs) family protein | 25.0 (2) |
| 116 | AT1G16760.1 | Protein kinase protein with adenine nucleotide alpha hydrolases-like domain | 23.0 (7) |
| 117 | AT1G79190.1 | ARM repeat superfamily protein | 23.0 (7) |
| 118 | AT5G37010.1 | unknown protein | 16.0 (3) |

The results in the list of protein families were obtained on the basis of a comparison of the masses of obtained peptides and their fragments with those of the TAIR10 database, using the MASCOT program in combination with the MScan software (http://proteom.ibb.waw.pl/mscan /index.html). 118 proteins were identified in the interaktome studied, of which 5 represented canonical Sm proteins (SmD1a, SmD1b, SmD3, SmD3b, SmB). Interestingly, there was a trend similar to the results obtained for mRNA identified by RIP-seq. A significant part of the cytoplasmic Sm interaction was the ribosome-related / translational proteins (eIF2 / IF5 translation initiators, RL10e ribosome structural proteins, RS1, RS3, OVA2 tNA RNase synthase), mitochondria (SHM1 and SHM2 methyltransferases, IDH isocitrate dehydrogenase, ABC ATM1 transporters) ) and plastids / photosynthesis (RuBisCO RCA asset, TKL1 transketolase). In addition, 2 families of mRNA binding proteins: Alba and ECT were identified.

**Supplementary Table 2 The cytoplasmic Sm proteins interactome. The results in the list of protein families.**

| locus | protein | score (# of unique peptides) |
|---|---|---|
| AT1G29250.1,AT2G34160.1 | Alba DNA/RNA-binding protein | 650.0 (4) |
| AT3G04620.1 | Alba DNA/RNA-binding protein | 650.0 (3) |
| AT4G19045.1,AT5G45550.1 | MOB kinase activator-like 1 (MOB1-LIKE) | 201.0 (2) |
| AT3G53510.1 | ABC-2 type transporter family protein (ABCG20) | 184.0 (4) |
| AT4G34200.1 | D-3-phosphoglycerate dehydrogenase 1 (PGDH1) | 163.0 (6) |
| AT1G76950.1 | Regulator of chromosome condensation (RCC1) family (PRAF1) | 151.0 (4) |
| AT3G55510.1 | Noc2p family (RBL) | 81.0 (1) |
| AT1G15690.1,AT1G15690.2 | Inorganic H pyrophosphatase family protein (AVP1, AVP-3) | 54.0 (1) |
| AT1G06590.1 | unknown protein | 51.0 (3) |
| AT3G48190.1 | PCD IN MALE GAMETOGENESIS 1 (PIG1) | 46.0 (15) |
| AT5G15070.1,AT5G15070.2 | Arabidopsis homolog protein of yeast VIP1 2 | 43.0 (3) |
| AT5G61150.1,AT5G61150.2 | VERNALIZATION INDEPENDENCE 4 (VIP4) | 28.0 (4) |
| AT3G07920.1 | Translation initiation factor IF2/IF5 | 258.0 (2) |
| AT5G22440.1,AT5G22440.2 | Ribosomal protein L1p/L10e family (RL10e) | 160.0 (4) |
| AT5G64140.1 | Ribosomal protein S28 (RPS28) | 110.0 (1) |
| AT5G49030.1,AT5G49030.2,AT5G49030.3 | tRNA synthetase class I (I, L, M and V) family protein (OVA2) | 87.0 (10) |
| AT1G12920.1,AT5G47880.1,AT5G47880.2 | Eukaryotic release factor 1 (ERF1) | 79.0 (4) |
| AT5G30510.1 | Ribosomal protein S1 (RPS1) | 70.0 (3) |
| AT1G50770.1 | Aminotransferase-like, plant mobile domain family protein | 68.0 (4) |
| ATMG00090.1 | ribosomal protein S3 (RS3) | 47.0 (2) |
| AT1G67120.1 | Midasin | 38.0 (18) |
| AT5G26780.1,AT5G26780.2,AT5G26780.3 | Serine hydroxymethyltransferase 2 (SHM2) | 1104.0 (7) |
| AT5G26860.1 | lon protease 1 (LON1) | 153.0 (4) |
| AT2G35120.1 | Single hybrid motif superfamily protein | 118.0 (2) |
| AT2G17130.1,AT2G17130.2,AT4G35260.1 | Isocitrate dehydrogenase subunit (IDH) | 111.0 (3) |
| AT3G53940.1 | Mitochondrial substrate carrier family protein | 42.0 (2) |
| AT4G28630.1 | ABC transporter of the mitochondrion 1 (ATM1) | 29.0 (1) |
| AT2G36250.1,AT2G36250.2 | Tubulin/FtsZ family protein (FTSZ2-1) | 439.0 (5) |
| AT3G52750.1 | Tubulin/FtsZ family protein (FTSZ2-2) | 326.0 (3) |
| AT2G45290.1 | Transketolase 2 (TKL2) | 290.0 (4) |
| AT1G26230.1 | Chaperonin 60 subunit beta 4 (CPN-60 beta 4) | 273.0 (2) |
| AT1G26230.2 | Chaperonin 60 subunit beta 4 (CPN-60 beta 4) | 273.0 (3) |
| AT4G00620.1 | EMBRYO DEFECTIVE 3127 (EMB3127) | 272.0 (2) |
| AT4G00600.1 | EMBRYO DEFECTIVE 3127 (EMB3127) | 272.0 (2) |
| AT2G04842.1 | EMBRYO DEFECTIVE 2761 (EMB2761) | 248.0 (4) |
| AT3G43190.1 | SUCROSE SYNTHASE 4 (SUS4) | 241.0 (5) |
| AT5G20830.1,AT5G20830.2 | SUCROSE SYNTHASE 1 (SUS1) | 241.0 (7) |
| AT5G49190.1 | SUCROSE SYNTHASE 2 (SUS2) | 224.0 (2) |
| AT4G02280.1 | SUCROSE SYNTHASE 3 (SUS3) | 224.0 (5) |
| AT2G39730.1,AT2G39730.2,AT2G39730.3 | Rubisco activase (RCA) | 145.0 (4) |
| AT3G54640.1 | Tryptophan synthase alpha chain (TSA1, TRP3) | 123.0 (3) |
| AT4G04730.1 | Unknown protein | 118.0 (4) |
| AT4G27070.1 | Tryptophan synthase beta-subunit 2 (TSB2) | 102.0 (3) |
| AT3G22960.1 | Pyruvate kinase family protein (PKP1, PKP-ALPHA) | 95.0 (7) |
| AT3G60750.1,AT3G60750.2 | Transketolase (TKL1) | 92.0 (4) |
| AT3G25820.2 | Terpene synthase-like sequence-1,8-cineole (TPS-CIN) | 79.0 (1) |
| AT1G59990.1 | EMBRYO DEFECTIVE 3108, DEAD-box ATP-dependent RNA h | 61.0 (3) |
| AT5G22450.1 | unknown protein | 57.0 (6) |
| AT3G54050.1,AT3G54050.2 | High cyclic electron flow 1 (HCEF1) | 34.0 (1) |
| AT1G67950.2 | RNA-binding (RRM/RBD/RNP motifs) family protein | 25.0 (2) |
